# CleLight: A scalable 3D histology pipeline for mapping neurodegenerative and psychiatric pathology in archival human brains

**DOI:** 10.64898/2025.12.10.693496

**Authors:** Tomàs Jordà-Siquier, Jules Scholler, Ivana Gantar, Héloïse Policet-Bétend, Laura Batti, Stephane Pages, Enikö Kövari, Christophe Lamy

## Abstract

Mesoscopic brain imaging, enabled by advances in tissue clearing, light-sheet microscopy, and large-scale image processing, allows detailed analysis of cellular and molecular architecture across whole neural circuits. However, applying these methods to postmortem human brain tissue is hindered by strong autofluorescence, high tissue density, and fixation-induced damage. We introduce CleLight, a light-enhanced clearing method that increases tissue transparency while quenching autofluorescence. Combined with complementary chemical treatments, CleLight supports multiplexed immunolabeling and high-resolution imaging of centimeter-thick, formalin-fixed, paraffin-embedded human brain sections. It is compatible with a wide range of antibodies and fluorescent dyes, enabling the visualization of physiological and pathological features across diverse CNS regions in healthy and diseased samples. CleLight offers a simple, robust, and scalable workflow for clearing, deep labeling, and volumetric imaging of human brain tissue. Its compatibility with conventional histology and archival material makes it well suited for organ-wide pathological studies in large patient cohorts and historical brain collections.

## Introduction

Understanding the cellular and molecular structure of the human brain is crucial for advancing neuroscience research. Neuroimaging techniques have significantly expanded our knowledge of the anatomy and function of the human brain in health and disease^1–3^. However, those methods lack the resolution and specificity required to define the brain’s cellular and molecular organization. Histological approaches, on the other hand, give access to neurobiological details in the brain tissue and are routinely used for neuropathological diagnosis^3,4^. They are classically constrained to 2D thin-sectioned samples which limits their ability to accurately capture complex structures like microvasculature and neuronal morphologies, sample heterogeneous cellular distributions and represent alterations distributed across brain regions.

Mesoscopic imaging methods, that aim at imaging organs in 3D with cellular resolution and molecular specificity, offer an opportunity to bridge the gap between neuroimaging and histology. They combine tissue clearing and labelling techniques with light sheet microscopy and advanced image analysis and visualization methods. Although tissue clearing was initially established for human tissue^52^, more recent implementations were focused of animal model systems^4,5^. Many of these methods use either hydrogels or aqueous solutions to preserve endogenous fluorescent proteins^6–8^. Although they have been tested on small human tissue samples (1-2 mm thick) ^9,10^, the use of denaturing agents like urea and SDS can affect the detection of certain epitopes by antibodies, thereby limiting their combination with immunolabeling. Furthermore, the frequent tissue expansion after tissue clearing can be problematic when scaling up to full human organ imaging.

Organic solvent-based clearing methods represent a valid strategy to clear and image larger organs in a short time^4,11,12^. They usually achieve a high level of clearing, are compatible with histological and immunostaining techniques, are simple to implement in a histopathology lab, can be automated, and enable the reuse of precious human tissue for classical or advanced thin-section imaging. This makes them ideal candidates for systematic use in neuropathology. However, clearing and staining archival human tissue poses significant challenges, including a high level of autofluorescence^13^, slow and partial antibody penetration in thick samples^12,14,15^, the detrimental effect on clearing and immunostaining of prolonged chemical fixation^14^, and a high tissue density due to myelin lipid density and sturdy molecules^16^, which necessitates the development of specialized tissue clearing methods.

To overcome those challenges, we developed CleLight, a solvent-based tissue clearing and staining method optimized for centimeter-thick formalin-fixed and paraffin-embedded postmortem human brain tissue sections. We combined light exposure with chemical treatments to enhance tissue discoloration, quench autofluorescence and provide superior transparency levels. Furthermore, we improved sample permeabilization and optimized the penetration and immunolabeling of a set of different antibodies and dyes that target various structures of the human central nervous system. With this protocol, samples could be reused for successive rounds of labeling, enabling highly multiplexed protein detection. Importantly, we could successfully process and immunostain archival samples preserved for up to 55 years in a historical brain tissue collection^17^. Although primarily developed to deal with human tissues, this method also improved tissue transparency of the mouse brain, indicating that it can be of more general use across species. Combined with our recent pAPRica petabyte image size reduction pipeline, CleLight provides a versatile and robust method to perform large scale 3D histopathological analysis of the human brain from current biobanks and historical brain collections.

## Results

### Light exposure quenches autofluorescence and increases brain tissue transparency

The age-dependent accumulation of autofluorescent and light absorbing molecules such as neuromelanin, oxidized flavins, lipofuscin, or collagen in the brain and the browning and hardening of brain tissue due to formalin fixation hampers clearing and imaging archival human brain samples^16,18^. Therefore, we assessed the efficacy and compatibility of quenching methods to reduce autofluorescence and enhance tissue transparency. Four different treatments—Sudan Black, copper sulfate, which were previously reported to successfully reduce autofluorescence^19^, as well as quadrol ammonium and light—were tested on archival brain tissue. Autofluorescence was assessed in unstained cleared samples illuminated at different wavelengths (488 nm, 555 nm and 647 nm). The intensity of autofluorescence was measured over ten orthogonal views along the illumination axis for each quenching method and an untreated control (Supplementary Fig. 1A, B). The treatment with sudan black had adverse effects on transparency, affecting light transmission through the tissue at different wavelengths. In contrast, copper sulfate effectively reduced autofluorescence but also had detrimental effect on tissue transparency. Next, we tested quadrol ammonium, a decolorizing reagent dissolving heme and other tissue pigments accumulating with age. It improved tissue transparency but did not significantly decrease autofluorescence. Eventually, we tried light exposure that proved to be the most effective treatment to reduce autofluorescence in all imaging channels, improving transparency (Fig. 1A, B, C; Supplementary Fig. 1A, B). We further investigated the effects of light exposure on brain samples from different patients with fixation times ranging between 6 months and 55 years (Fig. 1 C, D). Exposing cleared tissues to white light for 48 hours reliably quenched autofluorescence, regardless of the patient’s history or sample age (Fig. 1B, C; Supplementary Fig. 2A, B, 3A). In addition, we assessed the sample’s transparency by absorbance measurements on a spectrophotometer (Supplementary Fig. 1C). We confirmed that sudan black and copper sulfate led to a significant increase in light absorbance, limiting the light-sheet imaging depth. Quadrol ammonium and light exposure were the most effective treatments for decreasing absorbance, improving tissue transparency and enhancing imaging depth (Fig. 1E; Supplementary Fig. 1C). The increase in tissue transparency and quenching of autofluorescence after the light exposure was consistent across different samples (Fig. 1C,E) and was independent from light-induced temperature increase (Supplementary Fig. 3B) or any photobleaching effects caused by light-sheet fluorescence microscopic imaging (Supplementary Fig. 2D) or spectrophotometric measurements (Supplementary Fig. 2C). Overall, we concluded that light exposure and quadrol ammonium were the best treatments for rendering tissue transparent and quenching autofluorescence in archival brain tissue. We therefore decided to combine them for further experiments.

**Figure 1.**
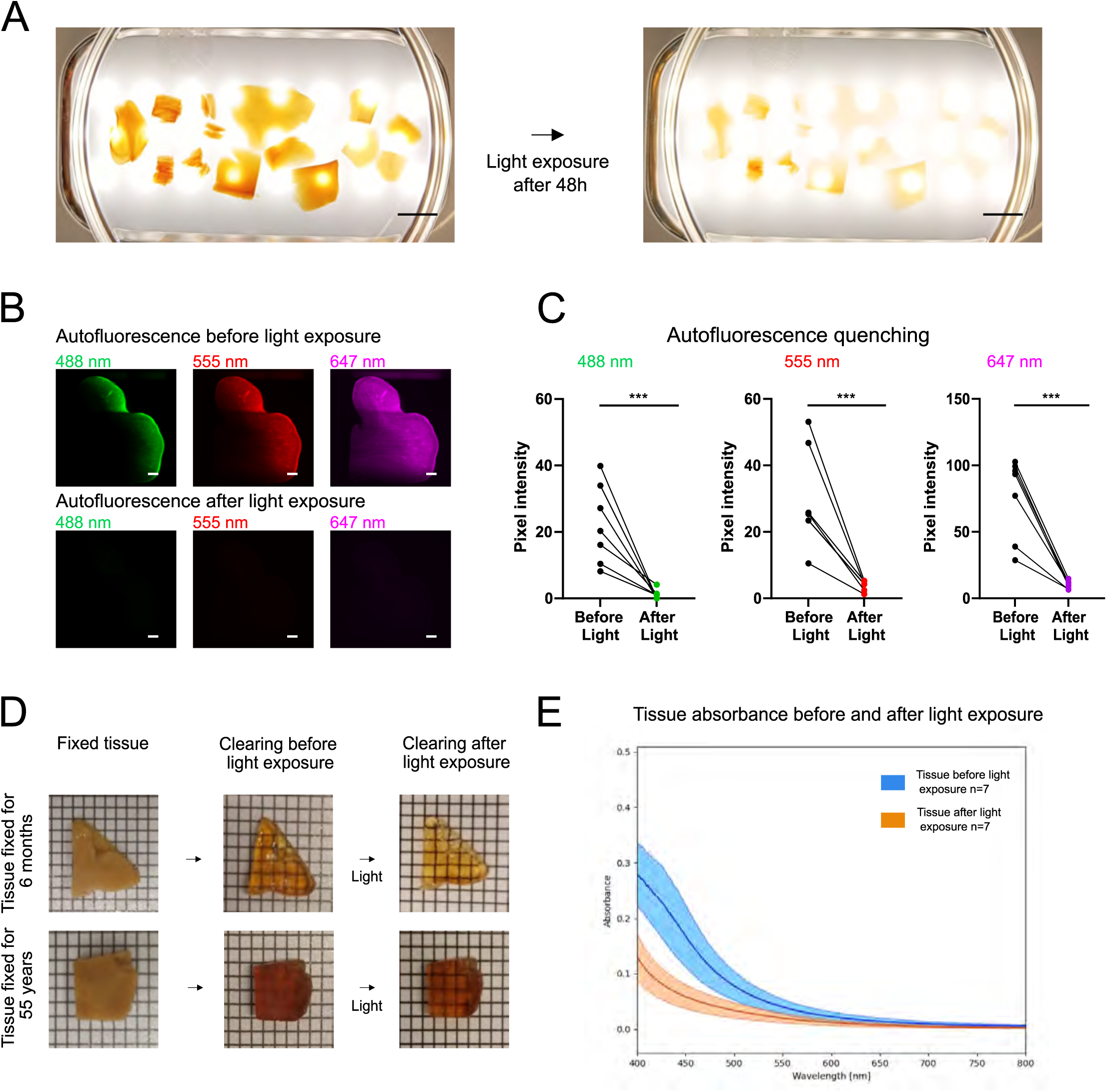
– Light exposure quenches autofluorescence and improves brain tissues transparency. **A)** Cleared human brain samples before and after exposure to white light for 48 hours. Scale bar = 1,5 cm. **B)** Light sheet images of autofluorescence at different wavelengths before and after 48h of white light exposure. Scale bars = 1 mm. **C)** Autofluorescence measurements in the different imaging channels before and after 48h white light exposure on 7 different brain samples coming from donors with different preservation times ranging from 6 months to 55 years. **D)** Images of brain samples with preservation times of 6 months and 55 years before clearing, after clearing and after light exposure (Grid: 2mm squares). **E)** Average light absorbance curve and standard deviation for the 7 brain samples before and after white light illumination. Absorbance measurements were done with a spectrophotometer.

### Improving immunostaining in large human brain samples

Immunostaining is necessary to visualize specific cellular and molecular structures in tissues. Performing this technique on thick post-mortem human brain samples is challenging due to the barriers of diffusion to large molecules, such as immunoglobulins, which result in suboptimal labeling of deeper regions. It is thus crucial to develop methods to improve the permeabilization of samples and facilitate the diffusion of antibodies in order to obtain a more homogeneous labeling in thick tissues. We explored several strategies, including the disruption of the extracellular matrix (ECM) by various methods and the use of smaller secondary antibodies, such as Fab fragment or nanobodies, to optimize this process. We first tested the loosening of the ECM either by enzymatic digestion with collagenase or hyaluronidase, or by a chemical treatment combining acetic acid and guanidine hydrochloride, as previously reported^12,20^. We incubated samples from the cerebral cortex of an Alzheimer disease (AD) patient overnight with either collagenase, hyaluronidase or acetic acid/guanidine hydrochloride under stringent conditions to prevent tissue degradation. Then we performed an immunolabeling for P-Tau, a protein involved in the formation of neurofibrillary tangles (NFT) in AD (Fig. 2A). In the control condition, the staining remained at the periphery of the samples, which was likely due to an accumulation of both the primary and fluorescently-labeled secondary antibodies at the tissue surface, preventing deep-tissue labeling and light penetration. Neither enzymatic treatments improved the labeling pattern as shown by the pixel intensity profile analyses (Fig. 2A, Supplementary Fig. 4A, B, C). In contrast, the chemical treatment combining acetic acid and guanidine hydrochloride resulted in a much more uniform staining across the entire sample volume, as shown by the intensity profile analysis, indicating improved diffusion of antibodies. (Fig. 2A, Supplementary Fig. 4D).

**Figure 2.**
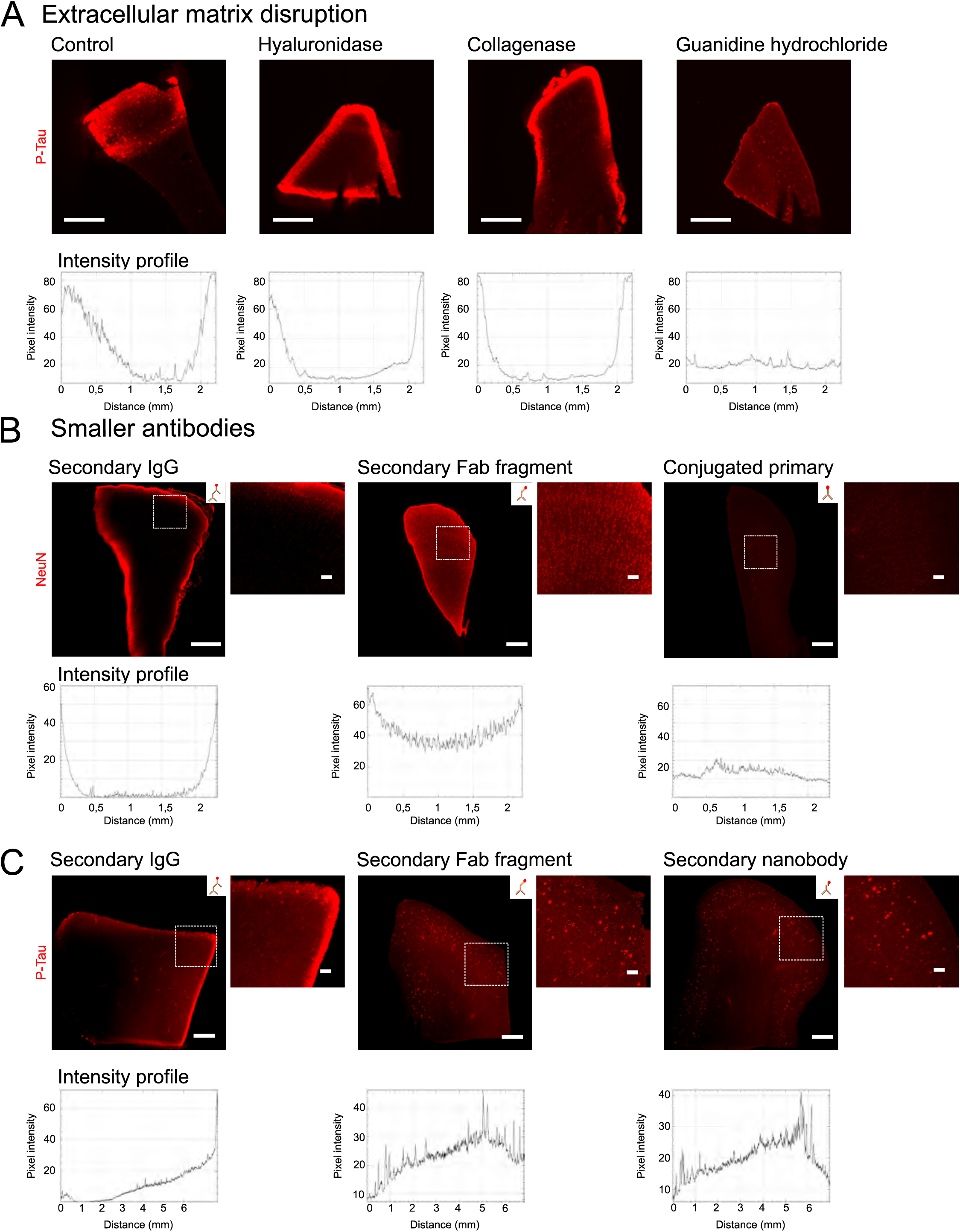
– Strategies to permeabilize human brain tissues. **A)** Efficacy of different ECM disruptors to improve antibody penetration in AD cortical tissues labeled with P-Tau, all from the same donor. **B)** Efficacy of secondary antibodies, Fab fragment secondaries and dye-conjugated primary antibodies to penetrate cortical samples from the same donor immunolabelled with the neuronal marker NeuN. **C)** Efficacy of secondary antibodies, Fab fragment secondaries and nanobodies to penetrate cortical AD samples immunolabelled with P-Tau present in NFT from the same donor and with pixel intensity profile distribution. Scale bars for main images = 1 mm and for zoomed acquisition insets = 100 µm. Lower rows indicate the relative pixel intensity profile of each corresponding image.

**Figure 3.**
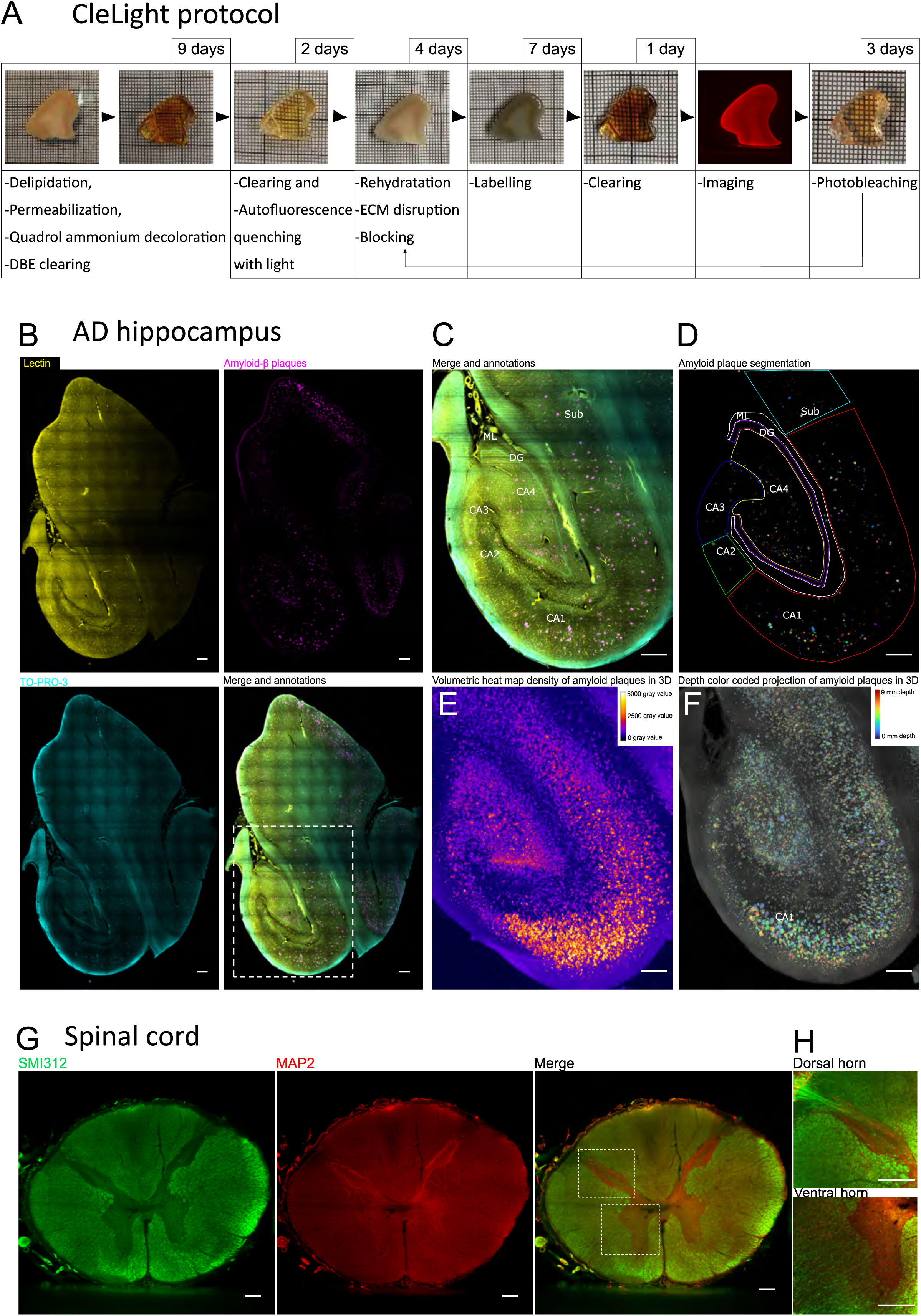
– CleLight enables high-resolution 3D reconstruction and analysis of large human brain samples. **A)** Graphic representation of the CleLight protocol with average processing times. **B)** Maximum intensity projection of the posterior human hippocampus (1cm thick section) from an AD donor labeled for vasculature (lectin), amyloid-β plaques and cell nuclei (TO-PRO-3). Scale bar= 1000 µm. **C)** Delineation of anatomical regions in the hippocampus. Scale bar = 1000 µm. **D)** 3D segmentation of amyloid-β plaques in different hippocampal regions. Scale bar = 1000 µm. **E)** Density of amyloid plaques in the human hippocampus represented as a heat map. Scale bar = 1000 µm. **F)** Depth of amyloid plaques represented as a color-coded projection. Scale bar = 1000 µm. **G)** Spinal cord from a control donor (2mm thick section) labeled for axons with neurofilament marker SMI312 and dendrites with MAP2. Scale bar = 500 µm. **H)** Maximum intensity projection of the dorsal and ventral horns of the spinal cord of a control donor. Scale bar = 500 µm.

We then tested a complementary strategy using antibodies smaller than full IgGs for immunolabeling, with the idea that they would diffuse better in thick tissue and therefore more easily access deeper tissue regions. We thus used either secondary antibodies made of IgG Fab fragments or camelid nanobodies, or directly fluorescently-labeled primary antibodies, to remove the need for secondary antibodies. We performed those tests on similarly sized pieces of cerebral cortex (1.5 x 2 x 2 mm).

Full IgG secondaries resulted in a highly inhomogeneous staining with neuronal marker NeuN (Fig. 2B, Supplementary Fig. 5A) or NFT marker P-Tau (Fig 2C, Supplementary Fig. 5B). In comparison, a one-step labeling procedure utilizing a complex of the primary antibody with a fluorescently-labeled Fab fragment secondary antibody formed prior to the incubation led to a much more homogeneous staining. A similar homogeneous staining was observed when using a secondary nanobody, to stain for pTau (Fig. 2C), or a directly fluorescently-labeled primary antibody, to stain for NeuN (Fig. 2B). Nevertheless, the signal obtained with the directly fluorescently-labeled primary antibody was weaker than when using Fab or nanobody-based secondaries, suggesting that the later compounds represent a better strategy for tissue labeling.

### Using CleLight for large scale studies of the human central nervous system histology and histopathology

Based on the previous experiments, we propose a solvent-based tissue clearing method optimized for large archival, formalin-fixed, brain samples combining: 1/ the quenching of autofluorescence and enhanced clearing by light exposure, 2/ the decolorization of samples by quadrol ammonium, 3/ the loosening of ECM with acetic acid and guanidine hydrochloride, 4/ the use of smaller secondary antibodies to improve immunolabeling homogeneity. This method, which we named CleLight, is outlined in Figure 3A.

We performed a series of experiments to demonstrate that this method can be used to label various physiological and pathological markers across the central nervous system. With CleLight, we successfully cleared the hippocampus, spinal cord, cerebellum and cerebral cortex, achieving consistent clearing across different patients and sample types (Supplementary Fig. 6). Dyes and antibodies validated with this protocol are listed in Supplementary Table 2. As an example, we co-labeled a centimeter-thick hippocampus sample from an AD patient with vascular marker lectin, amyloid-β plaques marker anbtibody, and the nuclear marker TO-PRO-3 (Fig. 3B). Hippocampal anatomical regions were delineated (Fig. 3C) before subsequent preprocessing with the pAPRica pipeline^21^, enabling the segmentation and 3D reconstruction of amyloid-β plaques (Fig. 3D). Those data were then displayed as volumetric density (Fig. 3E), and depth (Fig. 3F) of plaques. We then studied amyloid precursor protein (APP), another pathological protein directly linked to the etiology of AD, which is present in neuronal somas or extra cellular accumulations^22^. We successfully co-labeled APP with alpha smooth muscle actin, prominently present in arteries and the axonal protein beta III tubulin in the hippocampus, enabling us to study the deposition of APP in neurites and the vasculature (Supplementary Fig. 8).

We then investigated the suitability of our method to investigate AD pathology in the cerebral cortex. We successfully labeled pathological markers amyloid-β plaques and NFTs in cortical tissue using antibodies or chemical dyes such as congo red and Thioflavin S (Supplementary Fig. 7A, B, C, D). They were found in grey matter regions as well as in blood vessels, as expected. Furthermore, the co-immunolabeling of microglia marker Iba1 and amyloid-β allowed the visualization of neuroinflammation throughout the entire slice and around amyloid plaques (Fig. 4B, Supplementary Movie 3).

**Figure 4.**
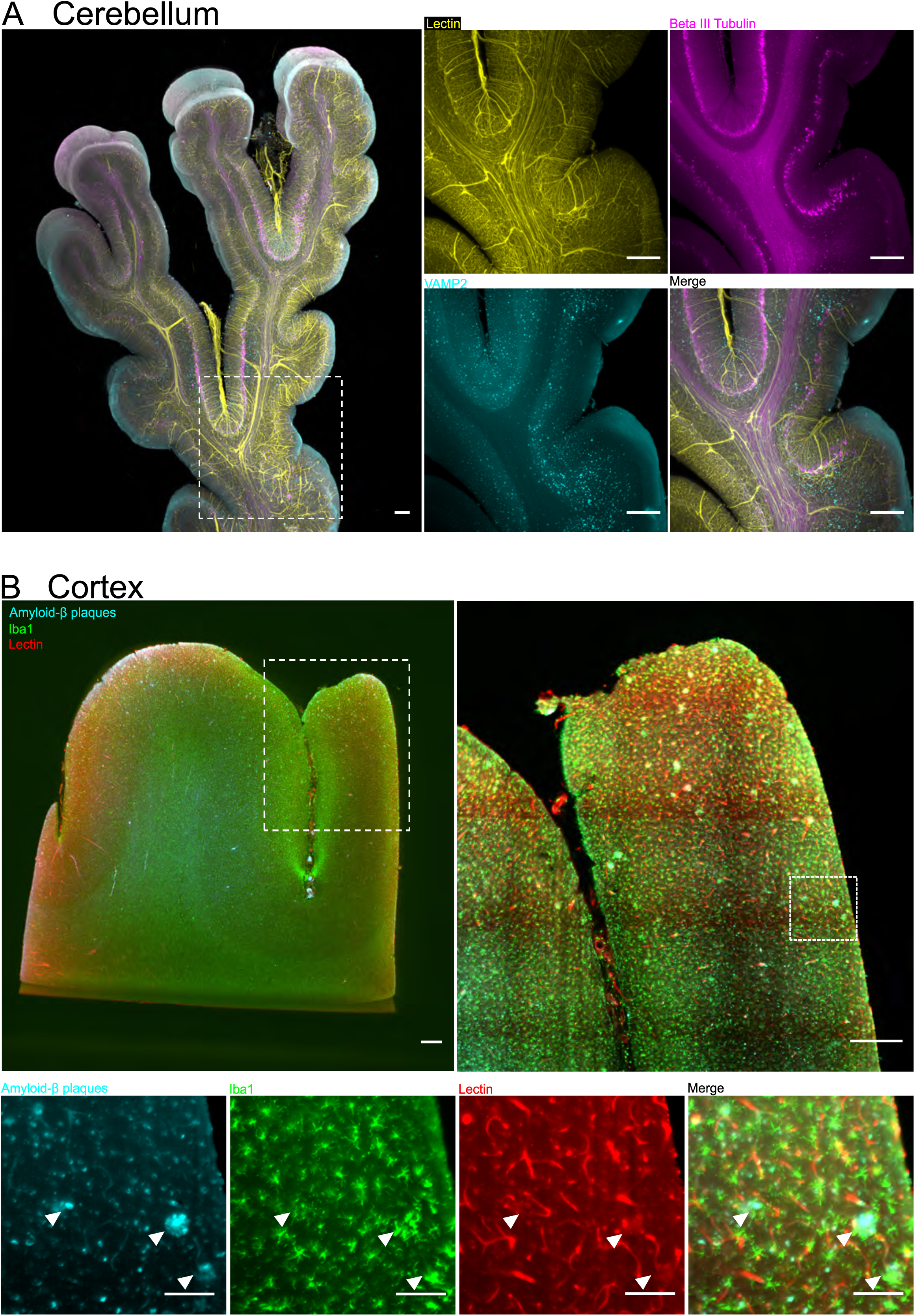
– **3D mapping of CNS structure and pathology with CleLight. A)** Maximum intensity projection of a cerebellar cortex section (4 mm thick) from a non-diseased donor labeled for vasculature (lectin), neurons (beta III tubulin) and synapses (VAMP2). Scale bar = 500 µm. **B)** Maximum intensity projection of a cerebral cortex section from an AD donor labeled for vasculature (Lectin), microglia (Iba1) and amyloid-β plaques. Scale bar = 500 µm

We assessed whether CleLight could help study the cytoarchitecture of non-pathological brain regions. To this aim, we labelled the vasculature and cytoskeletal markers present in axons and dendrites including beta III tubulin, anti-neurofilament SMI312, and MAP2 in the spinal cord (Fig. 3G, H, Supplementary Fig. 9D, Supplementary Movie 1), cerebellum (Fig. 4A, Supplementary Movie 2) and cortex (Fig. 4B, Supplementary Fig. 9B, Supplementary Movie 3). In the spinal cord, the labeling of dendrites with MAP2 delineated the grey matter whereas transversally cut myelinated axons were labeled by neurofilament SMI312 in the white matter (Fig. 3G, H, Supplementary Fig. 9D, E, Supplementary Movie 1). In the cerebellum, beta III tubulin defined the Purkinje cell layer as well as the axonal tracts in the white matter (Fig. 4A, Supplementary Movie 2). In the same tissue, vesicular synaptic marker Vamp2 identified the molecular and granular cell layers where synapses expressing this protein are located (Fig. 4A, Supplementary Movie 2). Identifying neurons and neuronal subpopulations is also key to delineating brain circuits. In cortical tissue, we used NeuN as a generic neuronal marker that enabled us to identify the different cortical layers (Supplementary Fig. 9C) and also specify the spatial distribution of pathological hallmarks such as amyloid plaques (Supplementary Fig. 7A, C, D, Supplementary Movie 4). Dopaminergic neuron’s marker tyrosine hydroxylase (TH) enabled the visualization of projecting neurites in the cerebellum (Supplementary Fig. 9A).

Because preclinical systems models are frequently used in neuroscience research, we sought to validate the CleLight protocol in non-human specimens. We processed full mouse brains with CleLight using various nuclear dyes and antibodies against densely expressed epitopes. CleLight led to a significant improvement in tissue clearing (Supplementary Fig. 10A) and enhanced diffusion of labeling molecules (Supplementary Fig. 10B, C) as compared to classical protocol iDISCO. Immunolabeling performed with CleLight resulted in stronger and more homogeneous labeling of Iba1 and histone H1 (Supplementary Fig. 10B). A similar improvement in staining homogeneity was observed with nuclear and vascular dyes TO-PRO-3 and lectin, respectively (Supplementary Fig. 10C). CleLight is thus suitable for both human and preclinical studies of the brain.

### Using CleLight to process archival formalin-fixed paraffin-embedded brain samples

Paraffin-embedding of formalin-fixed (FFPE) tissues is a standard technique in pathology departments prior to the preparation of thin sections for histopathological analysis. The remaining FFPE blocs are then stored in case further analysis is required, and sometimes indefinitely, constituting large tissue archives^17^. Because FFPE blocs are usually well preserved, those tissue archives are a precious resource for brain research on large cohorts of current or past conditions. We thus tested if CleLight can be applied to this type of material. To this end, we employed repeated thermal techniques to melt and remove the paraffin from the tissue, enabling subsequent processing with CleLight. Following this procedure, we cleared and stained full paraffin-embedded tissue blocks from various regions of the central nervous system, including the cortex, hippocampus, and mesencephalon, sourced from different patients (Supplementary Fig. 11A). Tissues were effectively cleared, indicating successful paraffin removal and the suitability of the CleLight protocol with this type of samples (Fig. 5A, B, C). Furthermore, protein epitopes were maintained, as evidenced by the successful immunostaining of samples. In samples from an AD patient, cortical tissue was well co-labeled with P-Tau together with lectin and TO-PRO-3, while the hippocampus was properly stained with congo-red, an amyloid fibril fluorescent dye labeling amyloid-β plaques (Fig. 5A, B). In addition, TH labeling of a mesencephalic section enabled us to evidence dopaminergic neurons and their projections of in various locations of the substantia nigra pars compacta, reticulata, and lateralis (Fig. 5C). Interestingly, the dopaminergic innervation delineated blood vessels, in register with its function if the regulation of vasoconstriction and vasodilation (Fig. 5C).

**Figure 5.**
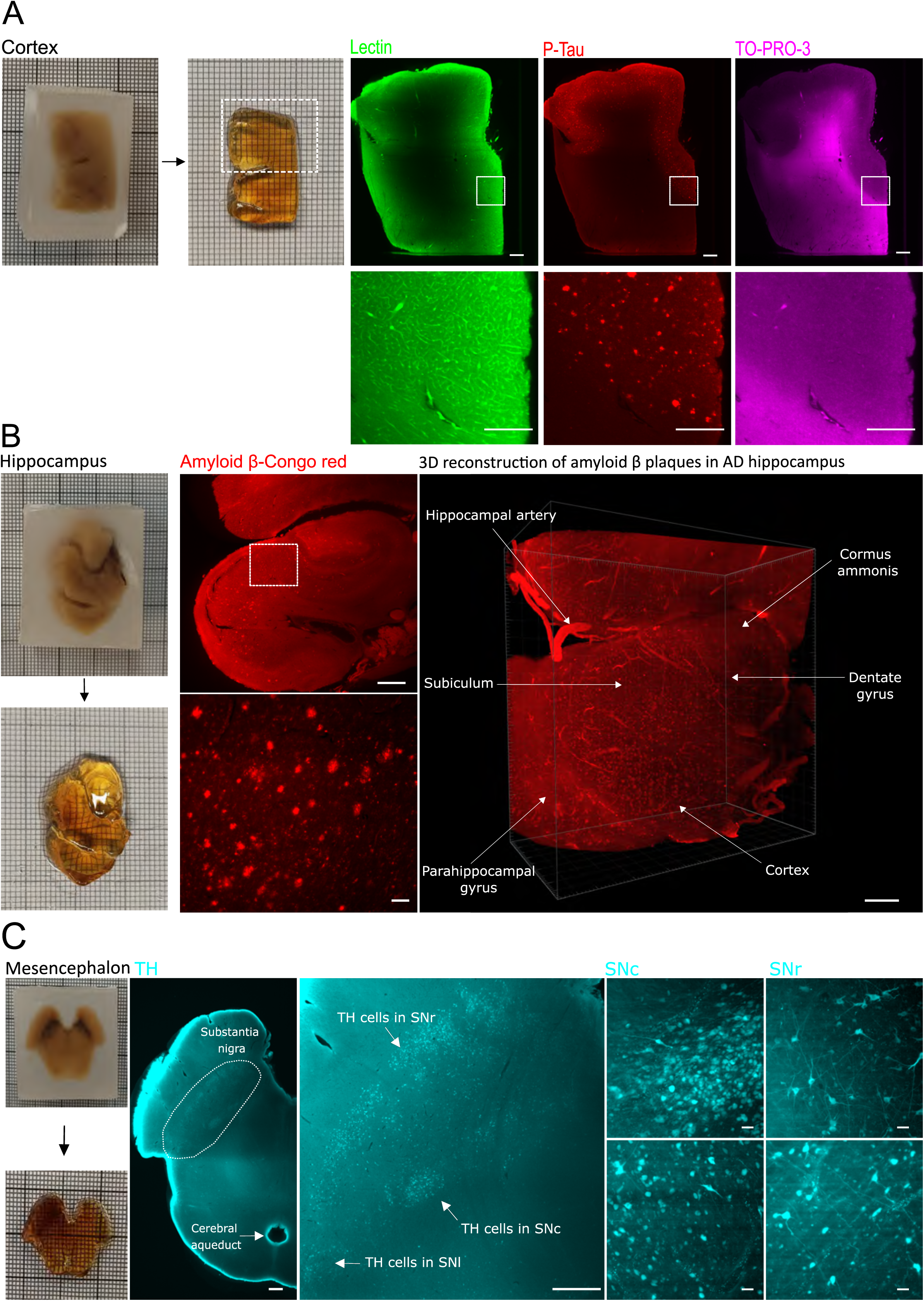
– Processing archival brain tissue with CleLight. **A)** Paraffin removal and clearing of an archival FFPE cerebral cortical bloc of from an AD donor labelled for vasculature (lectin), nuclei (TO-PRO-3) and P-Tau. Scale of macroscopic image = 1mm per grid square; Scale bars of low magnification images = 1 mm, high magnification images scale bars = 500 µm. **B)** Paraffin removal and clearing of an archival FFPE hippocampal block from an AD donor labeled for amyloid plaques with Congo red, shown at low and high magnification as single-plane images and as a 3D reconstruction. Scale of macroscopic image = 1mm per grid square; Scale bars for low magnification image and 3D reconstruction = 1 mm, high magnification image = 500 µm. **C)** Paraffin removal and clearing of an archival FFPE bloc of mesencephalon from a non-diseased donor immunolabeled for dopaminergic neurons (TH). Scale of macroscopic image = 1 mm per grid square; Scale bars for low magnification images = 1 mm, high-magnification images = 50 µm.

In addition to FFPE blocs, we processed formalin-fixed tissue preserved in formol. In cortical tissue, we observed that prolonged storage in formaldehyde led to the formation of crystalized deposits in the tissue, hindering light penetration and generating stripping artifacts during imaging (Supplementary Fig. 11B). We observed that applying a heat-induced epitope retrieval (HIER) protocol removed the crystals (Supplementary Fig. 11B), facilitating imaging in addition to improving immunolabeling by the re-exposure of protein epitopes.

Consequently, any samples stored for more than 2 years in formaldehyde underwent an antigen retrieval treatment. Those results demonstrate the general application of CleLight to both formalin-fixed and FFPE tissue archives, including to old archival samples such as in Fig. 4B and Supp 7D.

### Suitability of CleLight for 3D cyclic immunofluorescence (CycIF)

Cyclic immunofluorescence (CycIF), the ability to perform multiple rounds of immunolabeling on the same tissue enables highly multiplexed immunofluorescence imaging, which is essential to analyze precisely the complexity of normal and pathological brain tissue. In addition, it permits the extraction of the maximum amount of information from each precious human brain samples. We explored whether CycIF could be acheived with CleLight. A sample of hippocampus from an AD patient was labeled for alpha smooth muscle actin, APP C-terminal, and beta III tubulin (Fig. 6A, B, Supplementary Fig. 12A, B, C). After imaging, this sample was exposed to light for three days. Upon re-imaging, we detected no fluorescent signal in the sample, indicating successful photobleaching of the fluorophores (Supplementary Fig. 12B). Subsequently, we performed a second cycle of labeling for a new set of markers to assess the preservation of proteins. Lectin and fast green staining resulted in a strong fluorescent signal, indicating preservation of protein integrity and maintenance of tissue structure (Fig. 6C, D Supplementary Fig. 12C). This indicates that CleLight is compatible with CycIF procedures.

**Figure 6.**
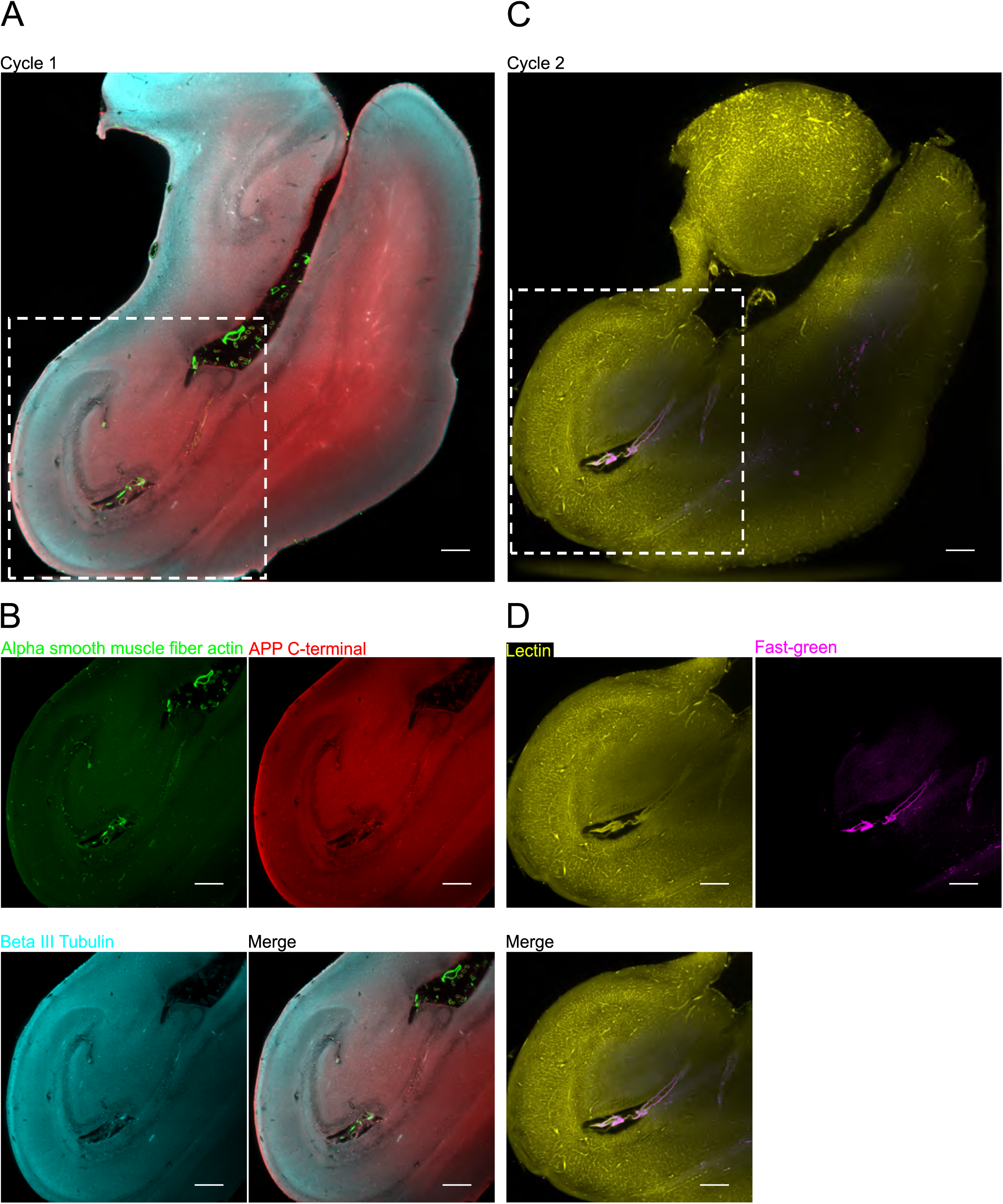
– Cyclic immunofluorescence with CleLight. **A)** Single plane microscopic image of an entire hippocampus from an AD patient labeled for blood vessels (alpha smooth muscle actin), pathology marker APP C-terminal and neurons (beta III tubulin). **B)** Higher magnification images of the previous sample. **C)** Single plane microscopic image of the same sample after photobleaching and a second round of labeling. The sample was relabeled for blood vessel (lectin) and collagen (fast green). **D)** Higher magnification images of the previous retained sample. Scale bars = 1 mm.

### Imaging whole human brain sections

Atlasing the entire human brain at the cellular level remains one of the biggest challenges in the field of neurosciences. To address this need, we integrated CleLight with pAPRica, our recently developed image processing pipeline^21^ to demonstrate its suitability to analyze full coronal brain slices from an AD patient.

One-centimeter-thick coronal brain slices, taken at the levels of the frontal lobe and the thalamus from an AD brain, were processed with CleLight, achieving complete transparency in both grey and white matter (Fig. 7A, Supplementary Movie 4). The frontal lobe level slice was subsequently labeled for amyloid-β plaques and lectin to visualize AD pathology and vasculature in the diseased brain. The image acquisition was performed with a theta light sheet microscope after sealing the slice in a chamber with DBE (Supplementary Fig. 13B,C)^23^. It covered 82 mm in width, 60mm in height, and 3mm in depth (301 planes), for a total of 41 x 30 tiles with a 5μm spacing resulting in a 10.9 TB dataset. The pAPRica pipeline reduced the dataset to 382.4 GB and enabled the stitching and merging of the acquired stacks, facilitating further visualization of (Fig. 7B). This integrated approach provides a comprehensive method for studying and understanding the intricate architecture of the human brain at large scale in health and disease.

**Figure 7.**
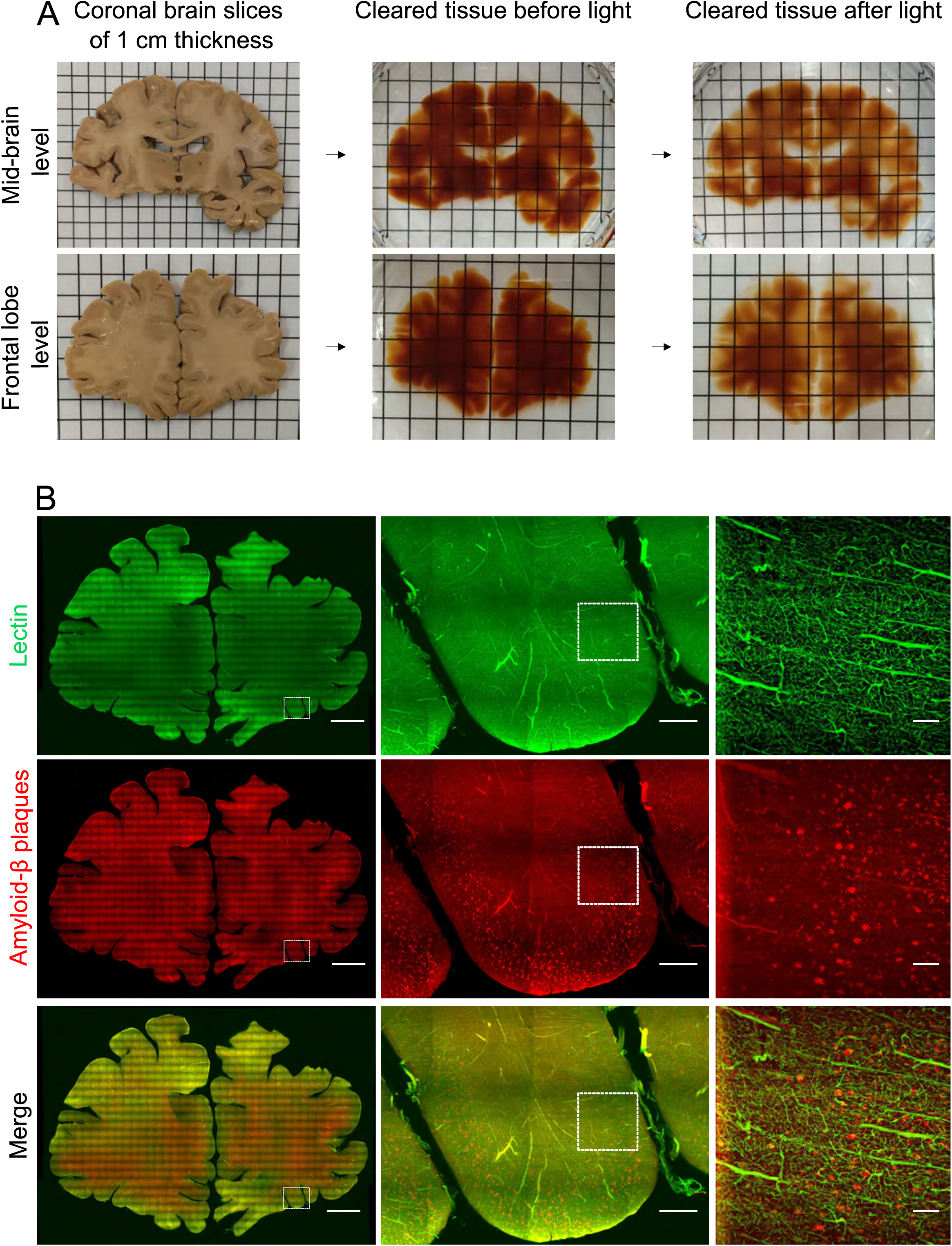
– Clearing of entire human brain slabs with CleLight. **A)** Centimeter-thick coronal brain slices from an AD donor at the level of the frontal and temporal cortices during the clearing by CleLight. Scale: 1 cm per grid square. **B)** Single planes images of a frontal cortex slice labeled for lectin and amyloid-β plaques. Scale bars for left images = 1 cm, middle images = 1 mm and right images = 100µm.

## Discussion

The emerging need for imaging the brain at cellular resolution at the whole organ level has led to the recent development of numerous tissues clearing methods. Many of these have been optimized for mouse models, enabling the clearing of entire brains and subsequent imaging with light-sheet fluorescent microscopy ^4,11^. Rodent models offer advantages for investigating cytoarchitecture such as minimal blood persistence and optimized fixation conditions thanks to perfusion fixation and the genetically-controlled expression of endogenous fluorescence proteins. In contrast, the human brain poses unique challenges for histological imaging. Age-related pigment accumulation leads to increased tissue opacity and autofluorescence^16,18^. In addition, prolonged preservation of tissue in formaldehyde leads to over-crosslinking of protein and masking of antigens^22,23^, which combined with high levels of myelination hampers clearing and prevents deep immunolabeling. To overcome these limitations, the volumetric study of the human brain requires optimized clearing methods that can effectively quench autofluorescence and improve clearing while enabling deep immunostaining throughout the entire sample thickness. In this study, we introduce CleLight, an optimized tissue clearing method designed to remove autofluorescence, increase tissue transparency and enhance labeling molecule diffusion in thick human brain slices.

Tissue clearing relies on reducing light scattering through refractive index (RI) matching and light absorption via pigment removal^13,24^. We started from the iDISCO protocol that uses methanol for dehydration, dichloromethane for delipidation and dibenzyl ether (DBE) for RI matching^11^. Although this protocol was successful in the mouse brain, it led to suboptimal clearing in larger archival human brain samples. We thought that this could be due to the accumulation of light absorbing and endogenous fluorescent molecules in brain samples from aged subjects. In addition, the iDISCO method, as many clearing protocols, employs hydrogen peroxide (H_2_O_2_) as an oxidative agent to degrade pigments^11,12,25^, which was not effective in fully eliminating light absorbing and autofluorescent molecules from archival human brain tissue^12,19^. We did not choose to apply more aggressive bleaching treatments as the prolonged exposure to high H_2_O_2_ concentrations can damage tissue structure and affect endogenous fluorescent protein emission^26^. We therefore explored alternative methods to increase tissue transparency and reduce autofluorescence. We first tested various autofluorescence quenching methods used in classical neurohistology^27^. We observed that conventional techniques like Sudan black and copper sulfate treatments, while effectively reducing autofluorescence^19^ had a detrimental impact on tissue transparency. They indeed enhanced light scattering and diminished light transmission, making them less suitable for microscopic imaging from thick tissue samples. We then tested the effect of quadrol ammonium which has been widely used as a decolorizing agent for clearing bone samples and dissolving blood heme^28^. We show that in the human brain quadrol ammonium improves clearing and reduces light absorbance without quenching autofluorescence.

Consequently, we assessed the efficiency of an exposure of precleared samples to white light as a way of reducing autofluorescence. Not only did this photobleaching approach effectively quenched autofluorescence, but it also decolorized and cleared tissues, all without the potential detrimental effects of chemical treatments on tissue integrity and protein epitopes. White light has been used as a conventional method to quench autofluorescence in various types of human tissues in 2D^29,30^. However, to our knowledge the application of light for tissue clearing in human samples has been relatively limited^15,31^. In this study, we demonstrated that the exposure of transparent archival tissue to light for 48 hours significantly reduced autofluorescence and enhanced tissue clearing, irrespective of the increase of sample temperature by the light source or any photobleaching effects induced by light-sheet fluorescent microscopy exposure. We postulated that light triggers the photodegradation of pigments such as lipofuscin, melanin, or heme^16,18,32^, thereby diminishing their light absorption capacity and thus tissue opacity^24^. As a result, the preincubation with quadrol and light exposure complement each other, significantly enhancing the optical clearing of human brain samples.

The age-related accumulation of insoluble lipids and macromolecules, such as collagen, in the extracellular matrix (ECM), poses a significant challenge for clearing and immunolabeling the human brain in 3D^12,14,33,34^. In addition, the density of certain antigens and their cross-linking by formaldehyde create an insoluble meshwork that prevent homogeneous 3D staining. However, different physical and chemical methods have been developed to enhance molecule penetration in tissues^7,14,20,35,36^. In this study, we hypothesized that combining different treatments would be essential to achieve a full labeling of archival brain tissue throughout its depth. We determined that the association of heat-induced antigen retrieval, disruption of the ECM and utilization of Fab antibody fragments led to homogenous immunostaining. For samples that were stored in formaldehyde for over two years, heat-induced antigen retrieval was employed break the covalent bonds formed by formaldehyde fixation and re-expose protein epitopes. We tested previously used methods to loosen the ECM including digestion with proteases or acetic acid combined with guanidine hydrochloride^35^. Enzymatic digestion with collagenase or hyaluronidase have efficiently been tested in hydrophilic clearing methods like CUBIC^20^. Nevertheless, those treatments proved inefficient with CleLight, in which the digestion and diffusion of enzymes was restricted to the surface of the sample and caused tissue degradation. In contrast, the acidic chemical treatment, known to efficiently break down collagen fibers and thereby loosening the ECM, allowed the deep penetration of dyes and antibodies with our method, in agreement with previous descriptions in entire human organs^35^.

In addition, we showed the interest of a one-step procedure for homogeneous deep immunolabeling. To this aim, we used either directly-conjugated primary antibodies or preformed complexes either labeled Fab fragments or nanobodies. Previous studies demonstrated the advantages of using nanobodies for the whole-body immunolabeling in Thy1-GFP mice^37^ and Fab fragments for the uniform immunolabeling of mouse brains^20^. All strategies successfully led to a deep uniform of densely expressed antigens such as NeuN, P-Tau, and neurofilament in both physiological and pathological conditions. Importantly, secondary Fab fragment antibodies and nanobodies yielded a better signal-to-noise ratio, compared to directly coupled antibodies, which lacked contrast due to a lower number of fluorescent molecules attached to the antibody. Overall, our findings confirm the critical role of smaller antibodies for deep labeling in human tissue clearing techniques.

Furthermore, we demonstrated that the use of light for photobleaching represents an accessible and cost-effective method for performing multiple rounds of staining and highly multiplex labeling that maximizes the utilization of human tissue. While this approach has been recently employed for quenching fluorescent probes in a 2D in situ hybridization study^30^, it has not yet been tested in 3D human brain samples. However, it is important to note that light does disrupt the antibody-epitope binding, meaning that the amount of specifically immunolabeled proteins is still limited by the number of antibody species used. Therefore, the exploration of alternative multiplexing methods^38,39^ or stripping buffers^40^ that remove antibodies without damaging the sample will need to be further investigated.

We showed that CleLight is an efficient method for performing tissue clearing and immunolabeling in a wide variety of both paraffin-embedded and formalin-fixed samples, even after long fixation times. Consequently, it offers multiple practical applications for research involving archival human brain samples. CleLight proved compatible with a wide variety of structural and pathological markers in different regions of the central nervous system. Consequently, we were able to visualize and segment the distribution of pathological hallmarks, like APP, amyloid plaques and NFTs, in combination with blood vessels or cell types like microglia (Iba1) and neurons (NeuN). Additionally, we successfully labeled specific neuronal subtypes such as dopaminergic (TH) cells, along with axonal (beta III tubulin, Neurofilament axonal marker SMI312) and dendritic (MAP2) markers, demonstrating the preservation of protein epitopes even after prolonged periods of formaldehyde fixation. Importantly, the diffusion of chemical dyes such as lectin, nuclear dyes, Congo red, or Thioflavin S was faster and more efficient than antibodies due to their smaller molecule size, though their specificity must always be controlled. Furthermore, in our experience, the labeling of high-density antigens like neurons or cytoskeletal proteins depended more on antibody concentration than incubation time. To improve deep immunolabeling in thicker tissues, antibody top-ups may be required.

Several tissue clearing methods have been applied to the clearing and labeling of human brain samples. Some of these techniques, such as CLARITY, SHIELD, CUBIC, and SWITCH, were initially developed for rodent brains and are not optimal for large brain sample sizes^6,7,33,41^. Other methods were specifically developed for the human brain. OPTIClear consisted in an adapted RI matching solution that could, in combination with delipidation or not, clear human brain tissues up to 2.5 mm thick. It was not compatible with all immunohistochemical staining and decreased the fluorescence of some dyes, which would be a limiting factor when imaging thicker samples^14^. The MASH protocols use similar low-cost and easy to implement delipidation and clearing steps as CleLight but do not have the light clearing step, which we showed significantly improves transparency and autofluorescence quenching. In addition, they achieve RI matching with Ethylcinnamate (Eci), which we found to be less efficient than DBE for clearing sample thicker than a few millimeters. Similarly, to MASH, we showed that fluorescent dyes could be combined with CleLight, including for staining blood vessels with fluorescently-tagged lectin. The MASH fluorescent staining methods are thus likely to be compatible with CleLight, provided they are applied after the light bleaching step, and might be used sequentially with intertwined light bleaching steps for multiplexing^42–44^.

The SHANEL protocol, another solvent-based method, with improved permeabilization based on the use of detergent CHAPS, was successfully used for clearing and immunostaining human brain samples. Still, as MASH, it lacked autofluorescence management steps, thus hindering its use as initially described for archival human specimens^35,45^. The SHORT protocol adopted the SWITCH tissue processing protocol combined with detergent-based delipidation and tiodiethanol (TDE) RI matching to clear human brain sections up to 500 µm. The SWITCH method is interesting for its ability to improve tissue preservation fixed by immersion, as usually done in pathology laboratories, and could be also tested with CleLight. The SHORT protocol determined that copper sulfate, sodium borohydride and H_2_O_2_, was the best autofluorescence reducing combination. Although we did not test the effect of sodium borohydride, we found that copper sulfate, while effective at reducing autofluorescence, also impeded light penetration. It thus remains to investigate how the SHORT would perform in thicker tissue samples. Noticeably, SHORT could be applied to FFPE samples with a similar deparaffinization method as CleLight, making it suitable for processing archival tissue blocs as well^19,33,46^. Similar to SHORT, SHARD employed an intra-tissue polymerization method, SHIELD, followed by detergent-based delipidation, for tissue clearing. This process enabled high temperature antigen retrieval and multiplexed immunolabeling, similarly to CleLight. Surprisingly, photobleaching did not significantly reduce the autofluorescent background with this method. Furthermore, it was only demonstrated with 200 µm sections and staining homogeneity did not seem optimal, questioning its suitability for multi-millimetric tissue samples^47^. The HIF-Clear method also aimed at handling human FFPE samples. Through optimized deparaffinization and antigen-retrieval-like high-temperature delipidation using Tween-20, instead of SDS, the authors managed to not only clear tissue but also to improve immunostaining. Interestingly, this method enabled multiple rounds of active immunostaining by consecutive antigen-retrieval treatments. However, although full mouse brains could be cleared, the maximum thickness documented in human tissue was 1 mm. The performance of this method on thicker human FFPE blocs has to be tested^48^.

Finally, the ELAST methodology, that creates an elastic hydrogel in tissue, achieved clearing and homogeneous immunostaining in centimeter-thick brain sections. Noticeably, this pipeline included a photobleaching-based autofluorescence quenching step. It is not shown if this step contributed to improving clearing as we showed in CleLight. This promising methodology, primarily designed for mapping the whole human brain, would also be useful for archival brain samples. Though it might be less straight forward to implement than solvent-based methods in the context of a pathology laboratory or smaller scale neuropathology project. In addition, unlike CleLight, the mELAST process cannot easily be reversed to embed tissue in paraffin for thin sectioning, thus limiting the reuse of precious clinical samples^15,49^. Inspired by, and building on, those previous methods, CleLight aims at providing an easy to implement method for neuropathological analysis.

Handling large datasets generated by full organ imaging represents another milestone that needs to be addressed for studying the human brain in its full complexity. To tackle this issue, we utilized our recently developed pAPRica pipeline^21^ to stitch, merge, segment, compress, and display an entire centimeter-thick human brain slab imaged with the ClearScope® light-sheet microscope. This enabled us to map blood vessels and amyloid plaques in a full brain hemisphere, establishing the fundamental steps for sample clearing, imaging, and data processing represents the initial groundwork for atlasing the full human brain in the near future. To facilitate this process, new generations of light-sheet fluorescent microscopes will need to be developed to allow for easy mounting and imaging of these samples^50^.

In summary, CleLight presents a straightforward, cost-effective method for optical clearing, autofluorescence quenching and deep immunostaining in human brain samples. This method facilitates the examination of recent and ancient archival brain tissue, enabling multiplexed immunolabeling for comprehensive histological analysis in 3D. The integration of CleLight with advanced light-sheet fluorescent microscopic imaging and automated image processing tools is a valuable tool for atlasing the cellular and molecular structure of the human brain, a key step for advancing our understating of brain anatomy and pathology. A bench protocol is provide online (https://lamylab.github.io/clelight/).

## Materials and methods

### Human brain samples

Formalin-fixed and paraffin-embedded brain tissue samples were obtained from the Geneva Brain Bank in accordance with the Ethics Committee of the Canton of Geneva (CCER n°2017-01937). Human brains from clinically and neuropathologically diagnosed control subjects and AD cases were taken during autopsy and immersion-fixed for at least one month in 10% formalin. The brains were then cut in cm-thick coronal sections from which smaller blocs where taken and embedded in paraffin to prepare histological sections. Both formalin-fixed and paraffin-embedded tissues were then stored at room temperature (RT). The CleLight protocol was tested on tissue samples from different regions of the nervous system, with different disease states and preservation times. Critical factors such as the postmortem delay, the age at death or, for AD patients, the Braak stage were recorded (Supplementary Table 1).

### Formalin-fixed tissue preparation and clearing

Formalin-fixed samples were removed from formalin and washed overnight in PBS 1x (Phosphate buffered saline P4417-100TAB Sigma Aldrich) with 0.2% v/v Tween-20 (Pan Reac applichem A4974) and 0.02% w/v Sodium azide (Sodium azide Fluka Biochemika 71289) at RT. Samples that were fixed for more than 5 years were subjected to a heat-induced antigen retrieval treatment in citrate buffer (10mM, pH 6.0) (Merck 1.00244.0500) with Tween-20 (0,2%) for 10 minutes at 110°C. Samples were then dehydrated with incubation in graded methanol (Fisher Chemical M/4000/17) series (methanol/water: 50%, 80%, 100%, 100%) for 1 hour each at room temperature. Samples were further incubated in a 66% dichloromethane (DCM) (Fisher Chemical D/1852/17)/ 33% methanol solution overnight at room temperature. Next day, brain tissue was washed twice for 30 minutes in 100% methanol and bleached with freshly prepared 5% *H*_2_*O*_2_ (Hydrogen peroxide 35% Thermo scientific CAS 7722-84-1) in 100% methanol at 4°C. The tissue was then rehydrated with serial baths of methanol/water at 80% and 50% followed by an incubation with PBS/ 0.3% Triton X-100 (T9284 Sigma-Aldrich), for 1 hour each at RT. Afterwards the tissue was permeabilized for 3 days at 37°C with a PBS solution containing 3% Triton X-100, 0.3M glycine (Pan Reac applichem A1067), 5% DMSO (41640 Sigma-Aldrich) and 10 *mg/mL* Saponin (Sigma-Aldrich S4521). After 2 washes in PBS with 0.2 % Tween-20, the tissue was incubated overnight at 37°C in a solution containing 25% v/v quadrol (N,N,N’, N’-Tetrakis(2-hydroxzpropyl)ethylenediamine 98% Sigma-Aldrich 122262-1L) and 5% m/v ammonium hydroxyde (Ammonium hydroxide solution 25% NH_3_ basis Sigma-Aldrich 30501-1L) in water to decolorize and dissolve the heme remaining in the tissue. Samples were again washed in PBS with 0.2% Tween-20 5 times for 30 minutes each at RT Samples were dehydrated in sequential methanol baths of 1 hour from 50 to 100% and left overnight in the last bath until complete dehydration. Finally, after a 3-hour incubation in 66% dichloromethane (DCM)/ 33% methanol at room temperature, and two 15-min washes in 100% DCM, samples were rendered clear with Dibenzyl ether (DBE, 33630 Sigma-Aldrich) overnight. The following day, samples in DBE were exposed for 48h to a white LED light (LED Aluminor Mika 10.079.101) at 10°C to reduce autofluorescence and improve the clearing. Samples were then rehydrated following the steps used to clear the tissue in reverse order until reaching 50% methanol. After 1 hour in 50% methanol, samples were left overnight in PBS / 0.2% Tween-20 for full rehydration. At this stage the ECM of the tissue was loosened by doing a first overnight incubation in 0,5M acetic acid at RT and a second incubation with PBS, 3% Triton, 0,05M CH3COONa, 4M Guanidine hydrochlorid (Roth 6069.3) at 37°C. Samples were then blocked for 2 days at 37°C in PBS, 0.2% sodium azide, 0.2% gelatin (VWR chemicals 24350.262). Brain tissue was then incubated with a primary antibody at 37°C for a week, followed by 5 washes in PBS, 0.2% Tween-20 over one day, and the incubated with a secondary antibody for another week. A second strategy for the labeling was co-incubated with a primary antibody and a Fab fragment secondary andtibody, which improved the staining penetration and shortened the staining time to one week. Primary antibodies and Fab fragment secondary were preincubated for 30 minutes at RT before adding it to the tissue. Due to the possible cross reactivity of the Fab-fragments secondaries, the labeling of each protein was done separately (Supplementary Table 2). All primary antibodies were diluted in a solution consisting of PBS, 0.2% Tween-20, 0,05% azide, 10mg/ml heparin (Braun 2022511) and 5% DMSO. When used, small molecule fluorescent dyes were incubated in PBS with a higher NaCl concentration (500mM). After the labeling, the tissue was washed in PBS/0.2% Tween for 1 day. Finally, samples were cleared with progressive dehydration steps in methanol, followed by a 3-hour incubation in 66% dichloromethane (DCM)/ 33% methanol, and two 15-min washes in 100% DCM before DBE immersion. After imaging of the samples, photobleaching could be done by using a white LED projector (15000 Im, 4000 K neutral white, Lumak Pro, Hornbach, Switzerland) for 5 to 7 days at 10°C. Samples were placed in a DBE-filled glass container immediately on top of the projector’s glass window. Tissue could be then rehydrated and relabeled a second round.

### Paraffin embedded tissue preparation and clearing

Paraffin embedded tissue blocks were first melted in an oven at 70°C before being incubated in in PBS / 0.2% Tween-20 at 70°C with continuous steering. In the first day of deparaffination the PBS / 0.2% Tween-20 solution was refreshed every hour while keeping the temperature constant at 70°C. The following days the buffer was refreshed twice a day until no micelles of paraffin could be found floating. The tissue was then dehydrated in a series of 1-hour ethanol (493511 Sigma-Aldrich) baths, (50%, 70%, 90% and 100% x2) at RT and a final overnight incubation in Xylene (214736 Sigma-Aldrich) at RT. Once deparaffinized, the tissue was again rehydrated with ethanol baths in the inverse order and a last bath in PBS Tween-20 at RT.

### Permeabilization enzymes

Where indicated, enzymatic digestion was used to permeabilize human brain tissue. Samples were digested overnight at 37°C with Collagenase-P (11213857001 Sigma-Aldrich, 1 mg/mL) in a carbonate buffer consisting of 50mM Na₂CO₃ (497-19-8 Sigma-Aldrich), 50mM NaHCO₃ (144-55-8 Sigma-Aldrich), 150mM NaCl and 25-100 µM EDTA, adjusted to PH=10. Another enzymatic reaction was hyaluronidase (H4272 Sigma-Aldrich, 3 mg/mL) in CAPSO buffer consisting of 10mM CAPSO (C2278 Sigma-Aldrich) and 150mM NaCl (207790010 Acros organics), adjusted to pH=10 and incubated overnight at 37°C.

### Autofluorescence and absorbance quantifications

Autofluorescence was measured from samples imaged with a single-sided light-sheet illumination and quantified by measuring the pixel intensity in ten different planes throughout the entire sample using Fiji. For each condition, the ten selected ROIs were close to the illumination side of the light-sheet. An average of the ten-pixel intensity values was then calculated and compared across different quenching conditions. On the same samples, absorbance was quantified at selected wavelength (400-800 nm) using a spectrophotometer (Biochrom Ultrospec™ 7000 PC UV-Vis Spectrophotometer). Samples were contained in cuvettes containing DBE and a negative control without any sample.

### Light-sheet microscopic imaging

#### MesoSPIM

Light-sheet microscopy was used for imaging human brain samples. Particularly, we used MesoSPIM^51^, a dual-sided illumination axially-swept light sheet microscope with a multiline laser combiner (405, 488, 561 and 647 nm, Toptica MLE). MesoSPIM is also provided with a detection path comprising an Olympus MVX-10 zoom macroscope with a 1× objective (Olympus MVPLAPO 1×), a filter wheel (Ludl 96A350), and a scientific CMOS (sCMOS) camera (Hamamatsu Orca Flash 4.0 V3). The excitation paths have galvo scanners for the generation of light-sheet and the reduction of shadow artifactt. Furthermore, the beam waist is scanned using electrically tunable lenses (ETL, Optotune EL-16-40-5D-TC-L), which are synchronized with the rolling shutter of the sCMOS camera. This axially scanned light-sheet mode (ASLM) allows for a uniform axial resolution across the field-of-view (FOV) of 5 μm. Image acquisition was done with a custom software written in Python. Z-stacks were acquired with a zoom set at 0.63X (pixel size 10.52 μm), 0.8X (pixel size 8.23 μm), 1X (pixel size 6.55 μm), 1.6X (pixel size 4.08 μm) or 2X (pixel size 3.26 μm), 3.2X (pixel size 2.03 μm) as 2048x2048 pixels images spaced by 3 or 5 μm. Excitation wavelength were at 488, 561 and 647 nm with respective emission filters 530/43, 593/40, LP664 (BrightLine HC, AHF). Single or dual light-sheet illumination was used depending on the thickness of the sample.

### COLM

Clarity Optimized Light-sheet Microscope (COLM) imaging was performed with one-sided illumination with scanned light sheets at 488, 561 or 647 nm. Emitted fluorescence was collected wit a 10X XLFLUOR4X / 0.28 N.A objective and imaged on an Orca-Flash 4.0 LT digital CMOS camera at 4 fps, in rolling shutter mode. A self-adaptive positioning system of the light sheets across z-stacks acquisition ensured optimal image quality over the whole sample thickness. Z-stacks were acquired at 3 μm spacing with a zoom set at 10x resulting in an in-plane pixel size of 0.6 μm (2048x2048 pixels).

### ClearScope

An open-top light sheet theta microscope with a large lateral travel range was used to image the 1 cm thick human brain slice (ClearScope, MBF Bioscience) (Supplementary Fig. 13 B, C)^23^. This microscope has a large travel range stage (150mm x 100mm x 12mm) and high working distance objectives (4X/0.28 WD 29 mm) allowing the imaging of large and thick human brain sections. The clearing of the brain tissue was initially achieved with DBE and then mounted in a chamber containing Ethyl cinnamate (W243000-1KG-K Sigma Aldrich). The microscope consisted in a double illumination light-sheet with 488, 561 and 647nm excitation channels. The emitted fluorescence was collected with a 4X objective (Olympus XLFLUOR4X, 0.28 N.A.), through an emission filter (ET 405/488/561/640 nm Laser Quad Band Set, Chroma) and imaged with an Orca-Flash 4.0 LT digital CMOS camera at 4 fps, in a rolling shutter mode. Calibration and aligning of the light sheet waist with the focal plane of the detection was done prior to the image acquisition to correct for the refractive index inhomogeneity in the z axis. Z-stacks with a spacing of 5 *µm* were acquired with the 4x objective, with a pixel size of 1.02 *µm* (2048x2048 pixels) and a 10% overlap between different tiles.

### Miltenyi Biotech Blaze UltraMicroscope

The full mouse brain was imaged with the Blaze UltraMicroscope from Miltenyi Biotech. The microscope consisted in a double illumination light-sheet that could image in 488, 561 and 647nm laser channels with a dynamic focus positioning. A big sample chamber allowed the microscope to image multiple brains. The fluorescence was collected with a 1.1X objective (N.A. 0.1), filtered and imaged with a 4.2 Megapixel sCMOS camera. Z-stacks were acquired with a spacing of 4.8 μm and a zoom set at 0.66-2.75X resulting in an in-plane pixel size of 6.5 μm (2048x2048 pixels).

### Image processing

Light-sheet datasets were processed to perform 3D reconstructions, object segmentation and volumetric analyses using Fiji or Imaris (v10). The pAPRica pipeline was applied to datasets exceeding 1 TB, acquired with COLM or Clearscope, to convert them to APR format and subsequently stitch them for enhanced visualization in Imaris.

### Statistical analysis

GraphPad Prism 10.3 was used to perform the statistical analyses. Normal distribution of the data was tested with a d’Agostino and Pearson tests. Data that were normally and non-normally distributed were compared using a two-tailed t-test and Mann-Whitney rank test, respectively. Two-way analysis of variance was used for analysis with two different factors and differences between groups were considered statistically significant when P< 0.05.

## Acknowledgements

This work was supported by grants from de Swiss National Science Foundation (n°219457, ArchiMed project) and the Wyss Center for Bio and Neuroengineering (Human Brain Mapping project) to C.L.

