## Supplemental Information for "CleLight: A scalable 3D histology pipeline for mapping neurodegenerative and psychiatric pathology in archival human brains"

### Supplementary Information

**Supplementary Figures**

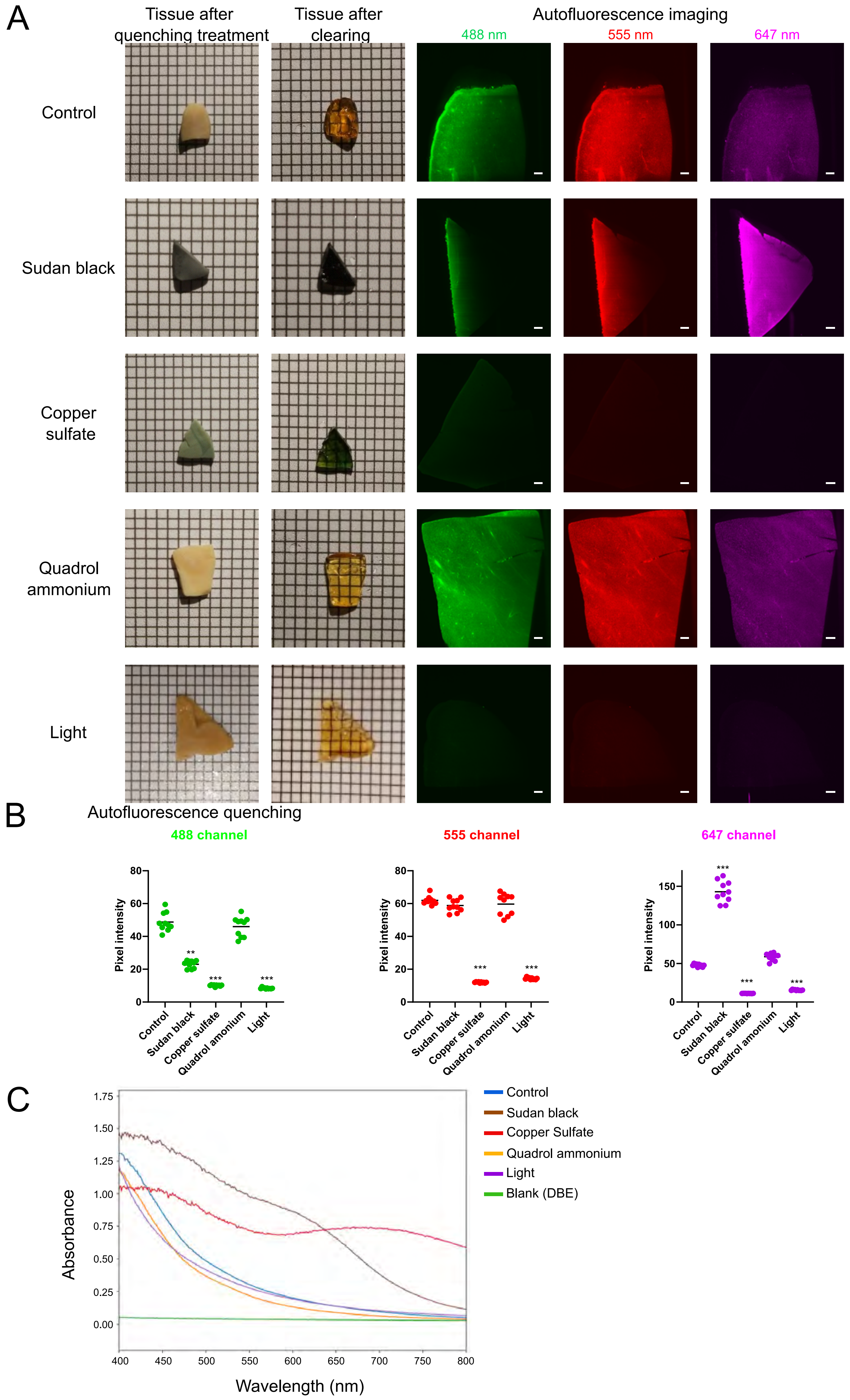
A
Tissue after quenching treatment
Tissue after clearing
Autofluorescence imaging
488 nm
555 nm
647 nm
Control
Sudan black
Copper sulfate
Quadrol ammonium
Light
B
Autofluorescence quenching
488 channel
555 channel
647 channel
Pixel intensity
Control
Sudan black
Copper sulfate
Quadrol amonium
Light
C
Absorbance
Wavelength (nm)
Control
Sudan black
Copper Sulfate
Quadrol ammonium
Light
Blank (DBE)

**Supplementary Figure 1 – Treatments for quenching autofluorescence in human brain tissues. A)** Cortical brain slices (2mm thick) from a control donor treated with different quenching methods before and after clearing (1mm per square). Endogenous fluorescence was imaged in different channels with a light-sheet microscope. Single plane images from the center of the sample are presented. **B)** Autofluorescence measurements at different wavelengths for the different treatment conditions. Average fluorescence intensity was measured on ten different planes along the z-axis and expressed as pixel intensity (A.U) on an 8-bit intensity scale (unpaired t-test $p<0.05$; $n=10$ planes for each group). **C)** Analyses of light absorbance for each treatment condition. Absorbance measurements were done in a spectrophotometer and expressed as a transmittance value.

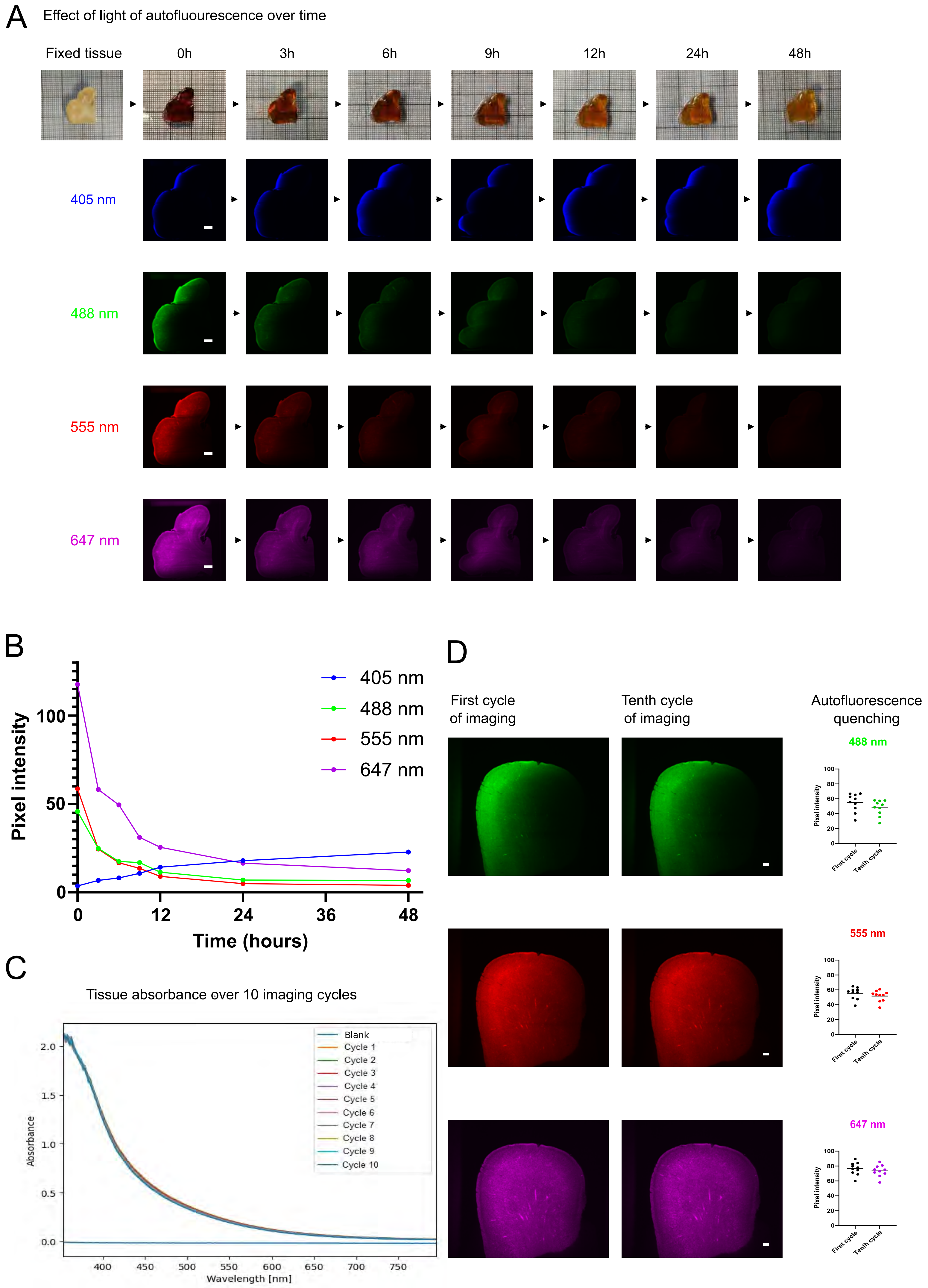
A
Effect of light of autofluourescence over time
Fixed tissue
0h
3h
6h
9h
12h
24h
48h
405 nm
488 nm
555 nm
647 nm
B
Pixel intensity
Time (hours)
405 nm
488 nm
555 nm
647 nm
C
Tissue absorbance over 10 imaging cycles
Absorbance
Wavelength [nm]
Blank
Cycle 1
Cycle 2
Cycle 3
Cycle 4
Cycle 5
Cycle 6
Cycle 7
Cycle 8
Cycle 9
Cycle 10
D
First cycle of imaging
Tenth cycle of imaging
Autofluorescence quenching
488 nm
555 nm
647 nm
Pixel intensity
First cycle
Tenth cycle

**Supplementary Figure 2 – Effect of white light exposure on autofluorescence and tissue clearing. A)** Light-sheet microscopy images of a cleared sample during exposure to light at four different wavelengths. Images were taken every 3 hours for 48 hours. **B)** Pixel intensity quantifications for each time point and channel. **C)** Absorbance measurements of tissue after ten cycles of the spectrophotometer. **D)** Light sheet images of a sample imaged iteratively for 10 cycles to assess the effect of imaging-induced photobleaching on autofluorescence. Images are shown of the 1$^{st}$ and 10$^{th}$ imaging cycle. Average fluorescence intensity was measured on ten different planes along the z-axis and expressed as pixel intensity on an 8-bit scale (A.U) (unpaired t-test $p<0.05$; $n=10$ planes for each group).

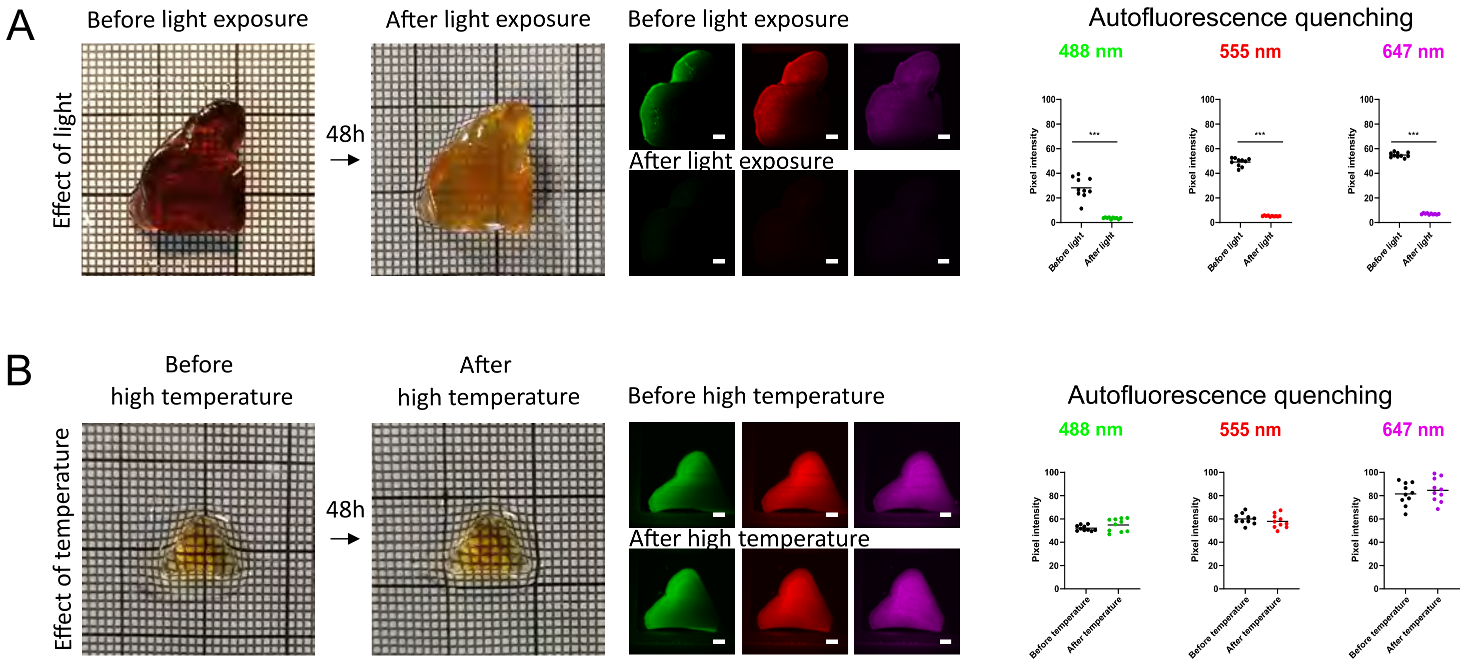


**Supplementary Figure 3 – Effect of temperature on autofluorescence. A)** Light sheet images of a cleared sample before and after 48h light exposure (unpaired t-test $p<0.05$; n=10 planes for each group). **B)** Light sheet images of a cleared sample before and after incubation at 37°C. (unpaired t-test $p<0.05$; n=10 planes for each group).

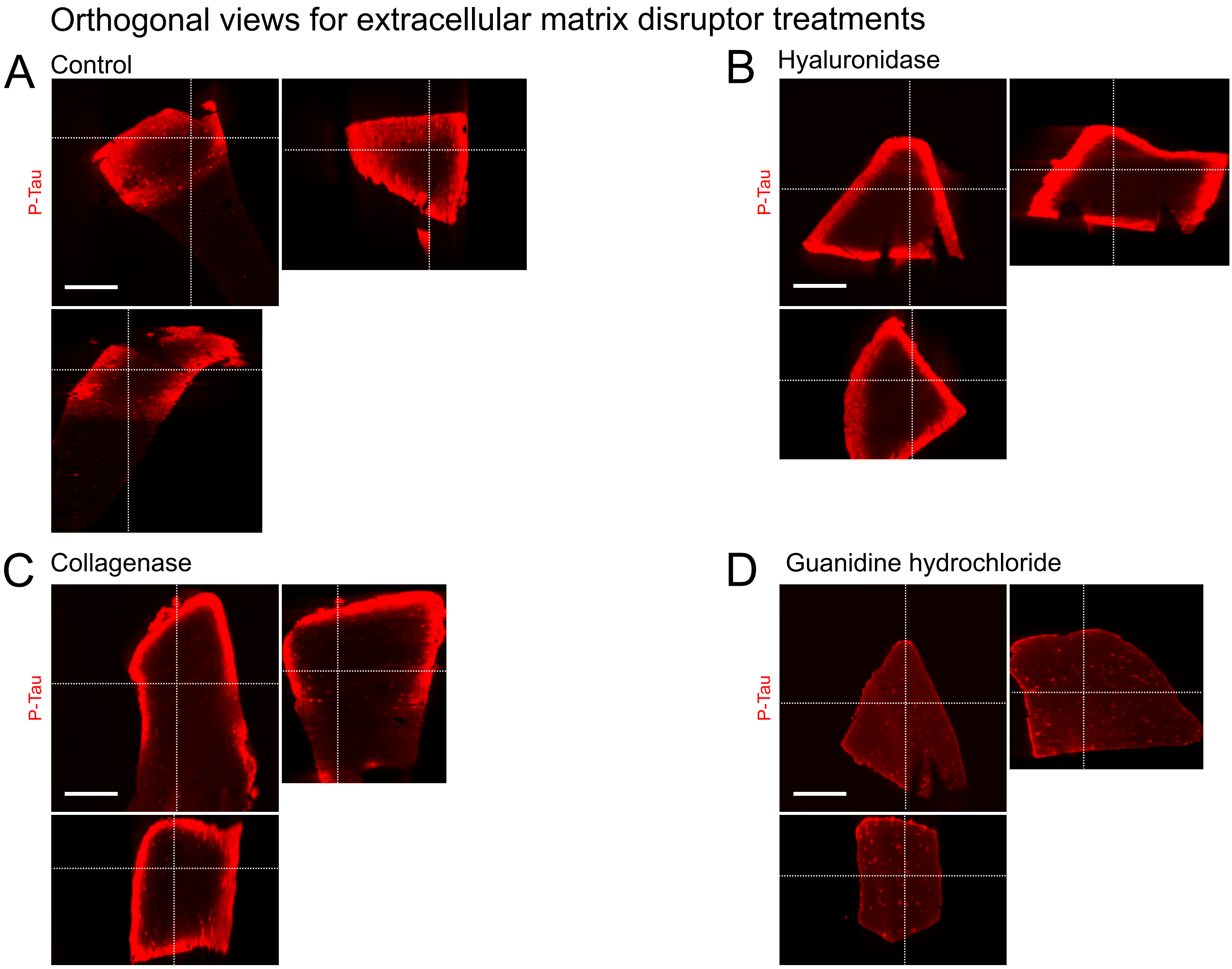


**Supplementary Figure 4 – Permeabilization and penetration of P-Tau antibody in different orthogonal views.** Penetration and labelling of P-Tau primary antibody with a secondary IgG antibody in different orthogonal views without treatment **(A)**, with a hyaluronidase treatment **(B)**, collagenase treatment **(C)** and guanidine hydrochloride treatment **(D)**. Scale bars = 500µm

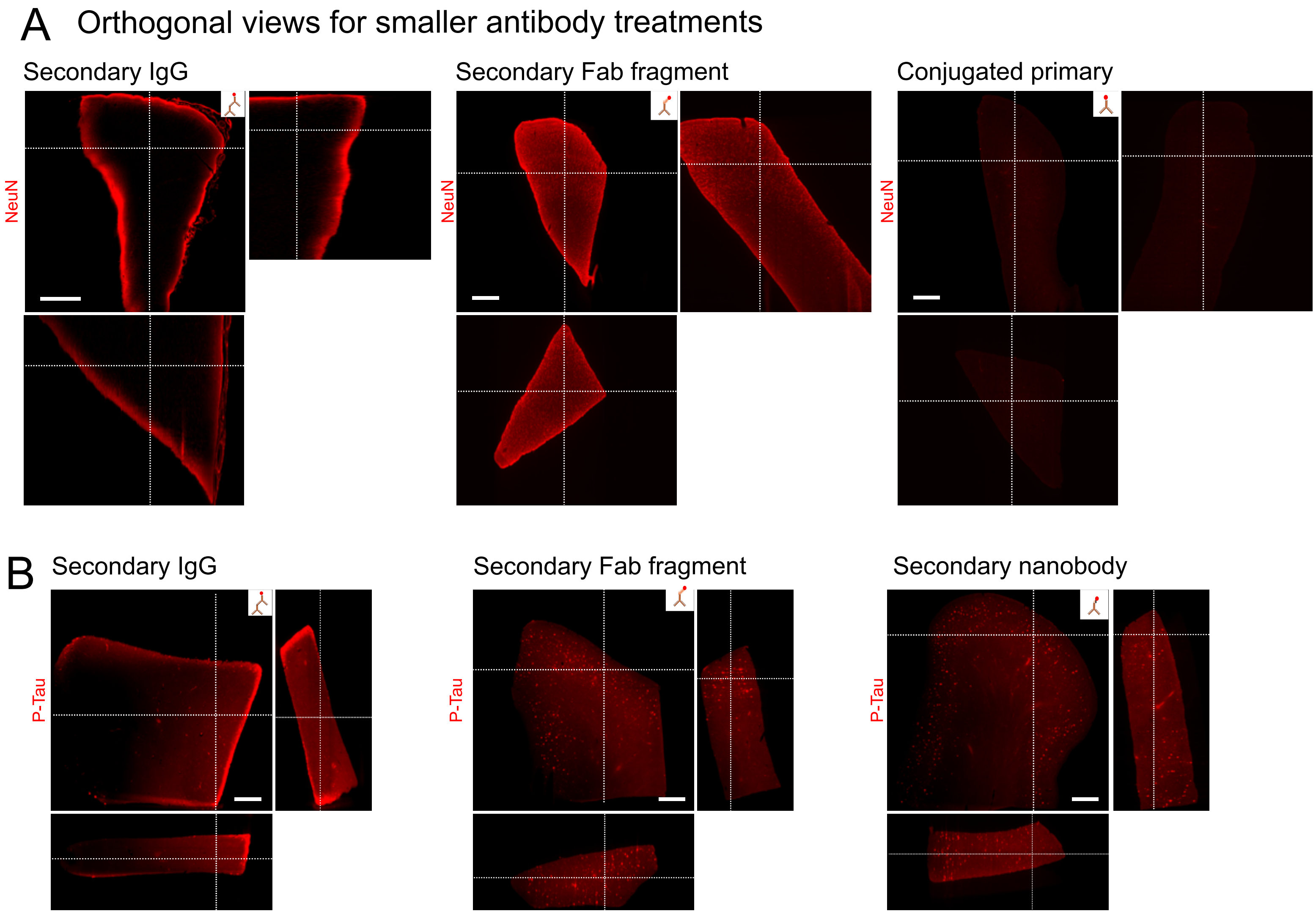


**Supplementary Figure 5 – Permeabilization and penetration of NeuN antibody in different orthogonal views. A)** Penetration and labeling of smaller antibodies for NeuN in different orthogonal views. Scale bar = 1mm. **B)** Penetration of smaller antibodies for P-Tau in different orthogonal views. Scale bars = 1mm

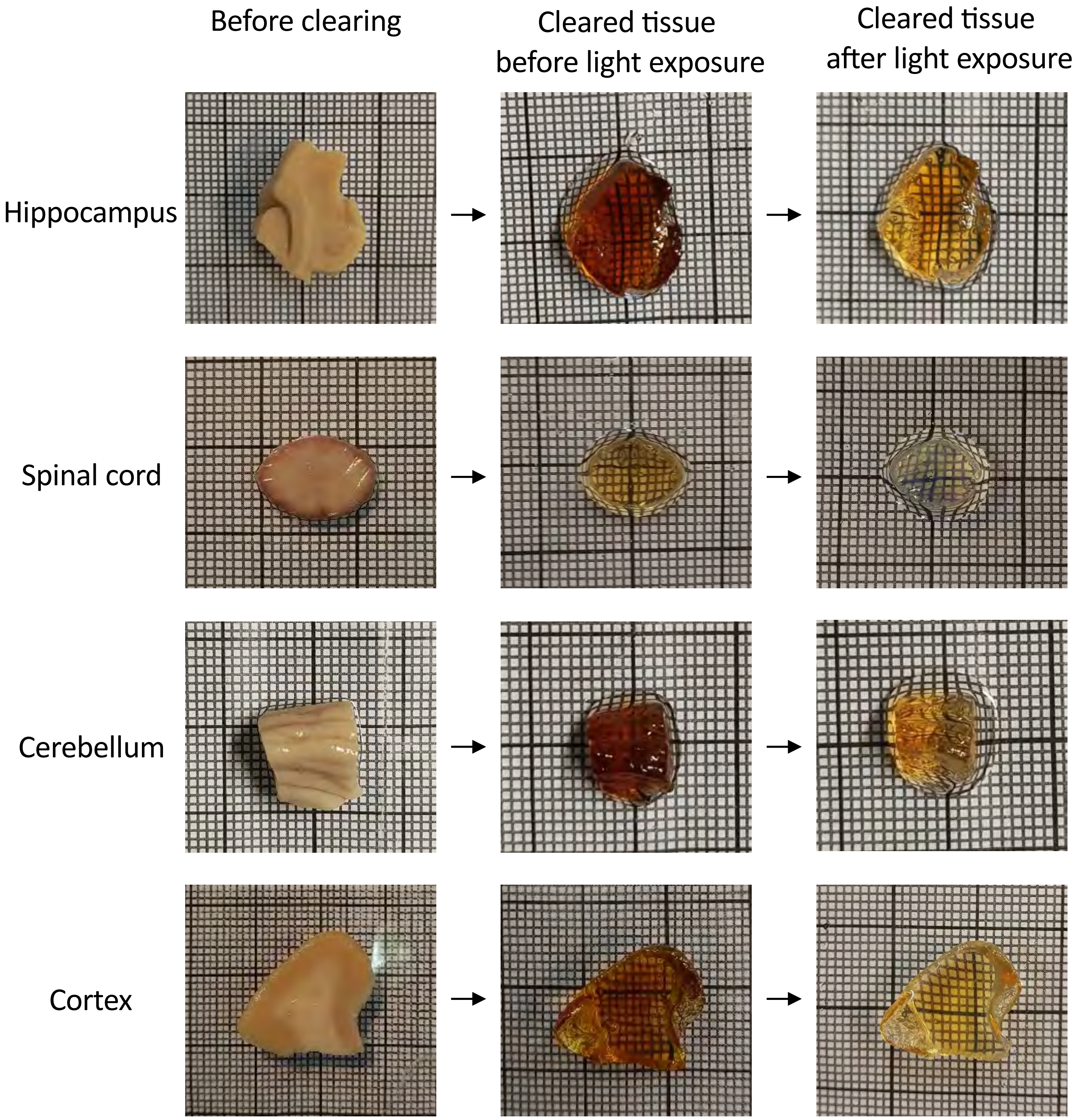


**Supplementary Figure 6 – Effect of light exposure in different regions of the central nervous system.** Effect of clearing with light in different parts of the central nervous system including hippocampus, spinal cord, cerebellum and Cortex (grid: 1mm per square).

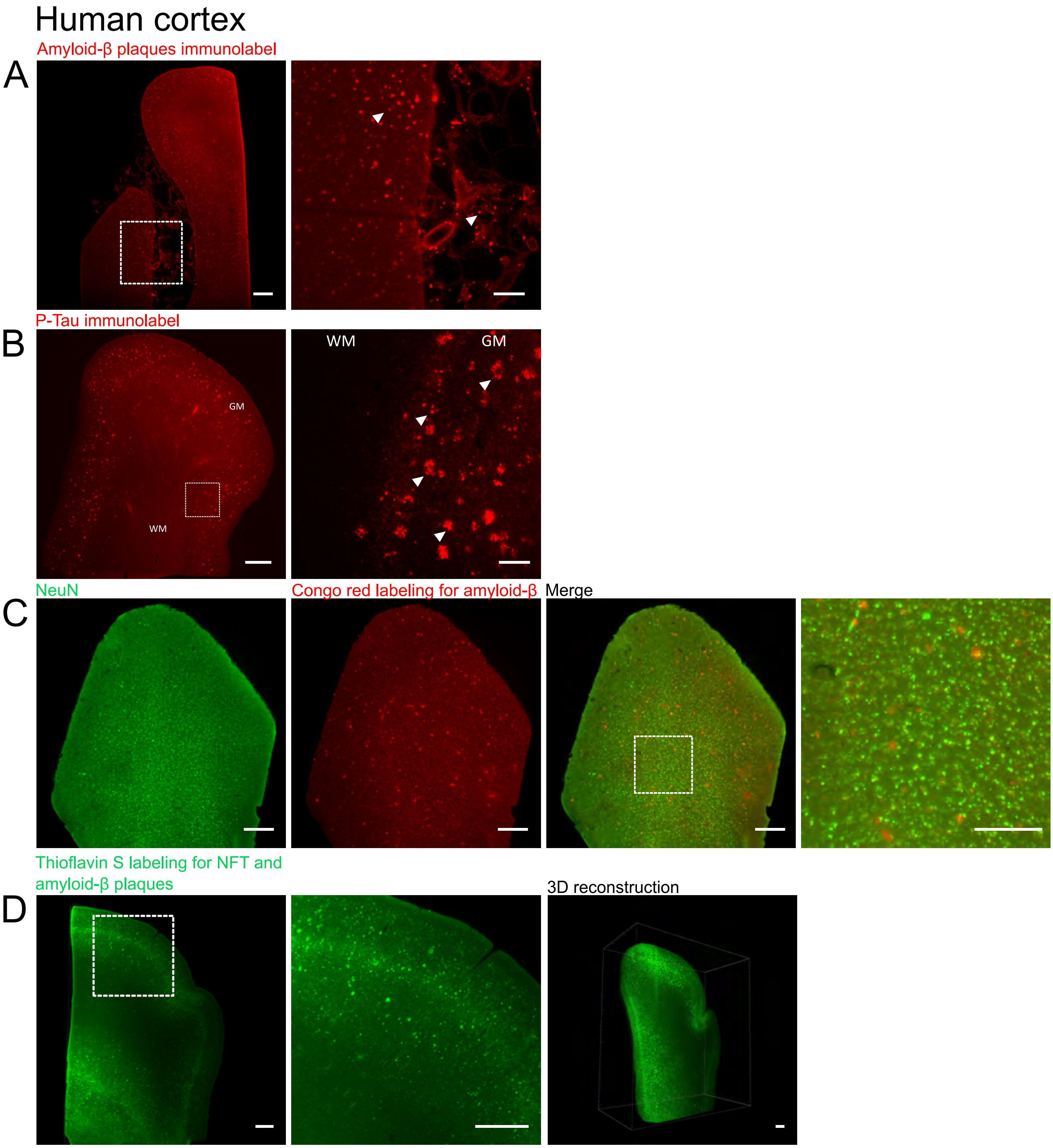


**Supplementary Figure 7 – CleLight enables 3D reconstruction and processing of large human brain samples. A)** Frontal human cortex from an AD donor patient (1cm thick) immunolabeled with amyloid-β plaques. Scale bar = 1000μm. Scale bar zoom = 500μm. **B)** Same AD donor patient and region immunolabeled with P-Tau present in NFT. Scale bar = 1000μm. Scale bar zoom = 500μm. **C)** Same AD donor patient and region labeled with NeuN and Congo red to visualize neurons and amyloid β plaques. Scale bar = 500μm. Scale bar zoom= 250μm. **D)** Archival brain tissue from an AD donor patient labeled with Thioflavin-S for amyloid-β plaques and NFT. Scale bar = 1000μm. Scale bar zoom = 1000μm.

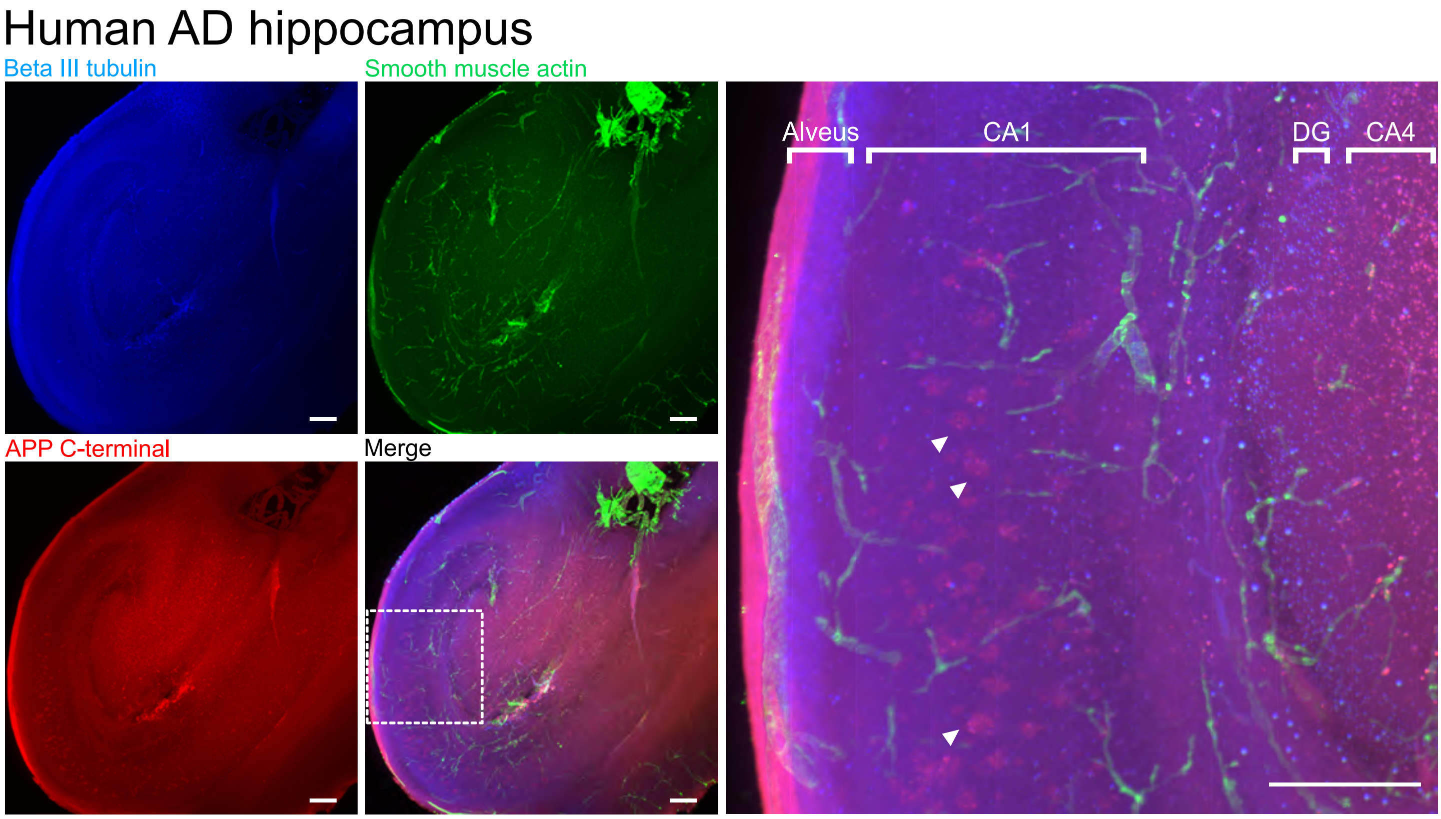


**Supplementary Figure 8 – CleLight enables 3D reconstruction and processing of large human brain samples.** Z-projection of a human AD hippocampus immunolabeled with Beta III tubulin, alpha smooth muscle fiber actin and APP C-terminal present in accumulations (arrow heads). Scale bar= 500µm. Scale bar zoom= 500µm.

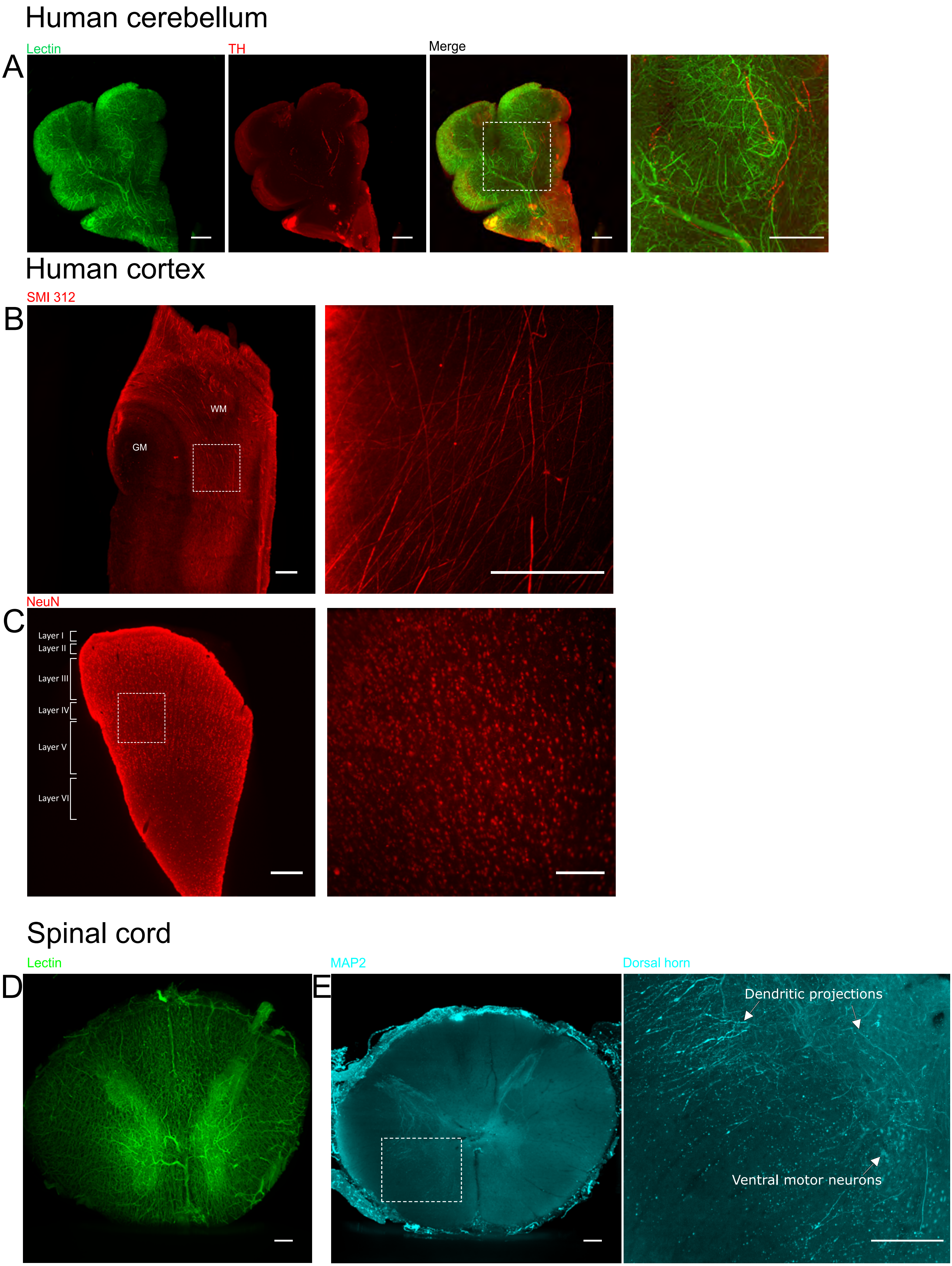
Human cerebellum
Lectin
TH
Merge
A
Human cortex
SMI 312
B
WM
GM
NeuN
C
Layer I
Layer II
Layer III
Layer IV
Layer V
Layer VI
Spinal cord
Lectin
MAP2
Dorsal horn
D
E
Dendritic projections
Ventral motor neurons

**Supplementary Figure 9 – CleLight enables 3D reconstruction and processing of large human brain samples.** A) Z-projection of the human cerebellum labeled with lectin and dopaminergic projections labeled with tyrosine hydroxylase. Scale bar= 500µm. Scale bar zoom= 500µm. B) Z-projection of cortical tissue labeled with the axonal marker for neurofilament SMI312 present in white matter (WM). Scale bar= 500µm. Scale bar zoom= 500µm.  C) Cortical layers identified in an immunolabeling of NeuN D) Z-projection of a spinal cord's vascularization labeled with lectin. Scale bar= 500µm. E) Z-projection of a spinal cord's dendrite innervation. Scale bar= 500µm. Scale bar zoom= 500µm.

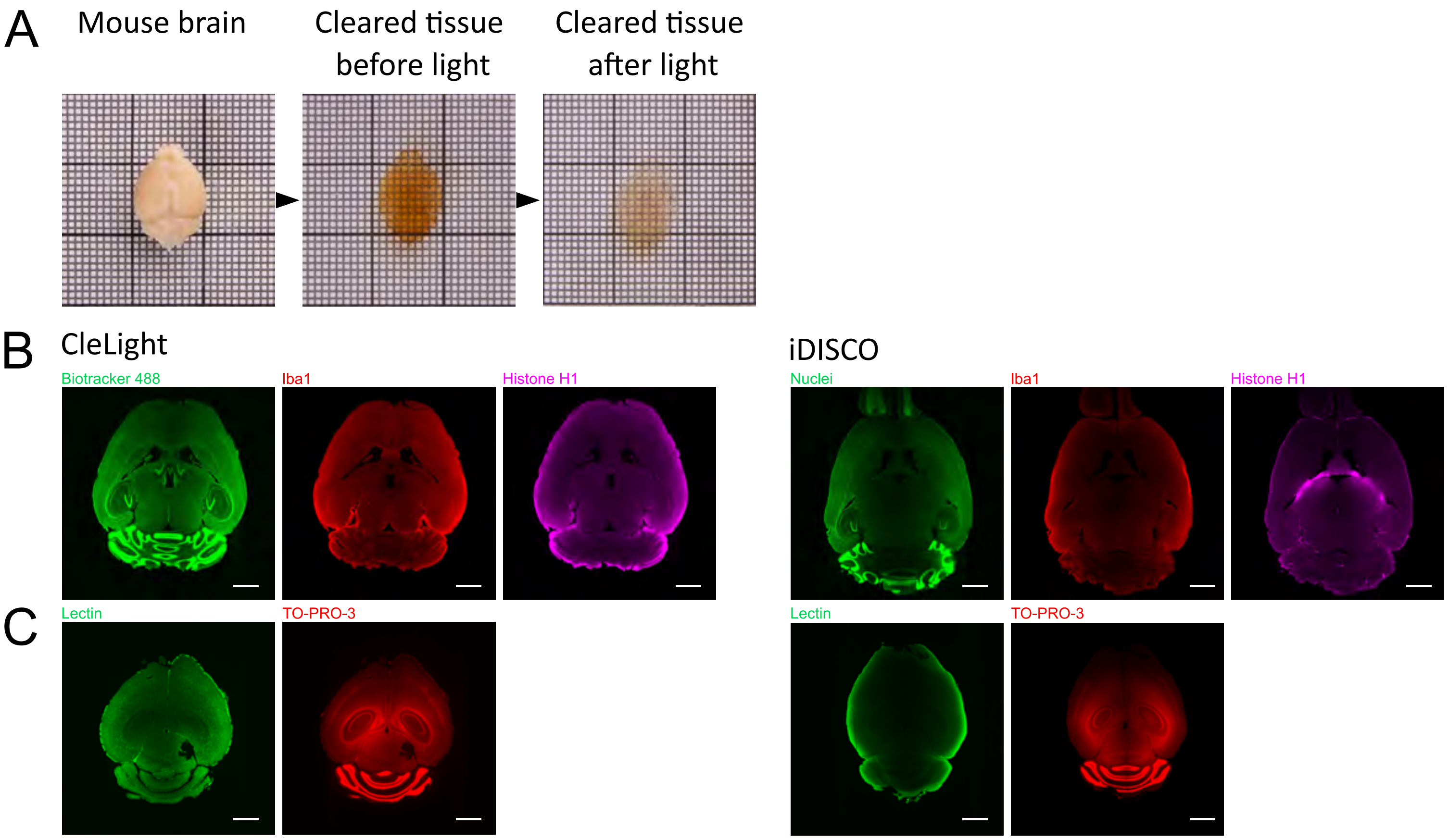


**Supplementary Figure 10 – CleLight enables 3D reconstruction and processing of the mouse brain. A)** Mouse brain processed with CleLight after light exposure (1mm per square). **B)** Diffusion comparison between CleLight and iDISCO after performing labelings with biotraker 488 (cell nuclei dye), microglia and Histone H1. Scale bar= 2mm. **C)** Diffusion comparison between CleLight and iDISCO for chemical dyes to label cell nuclei (TO-PRO-3) and vasculature (lectin). Scale bar= 500µm.

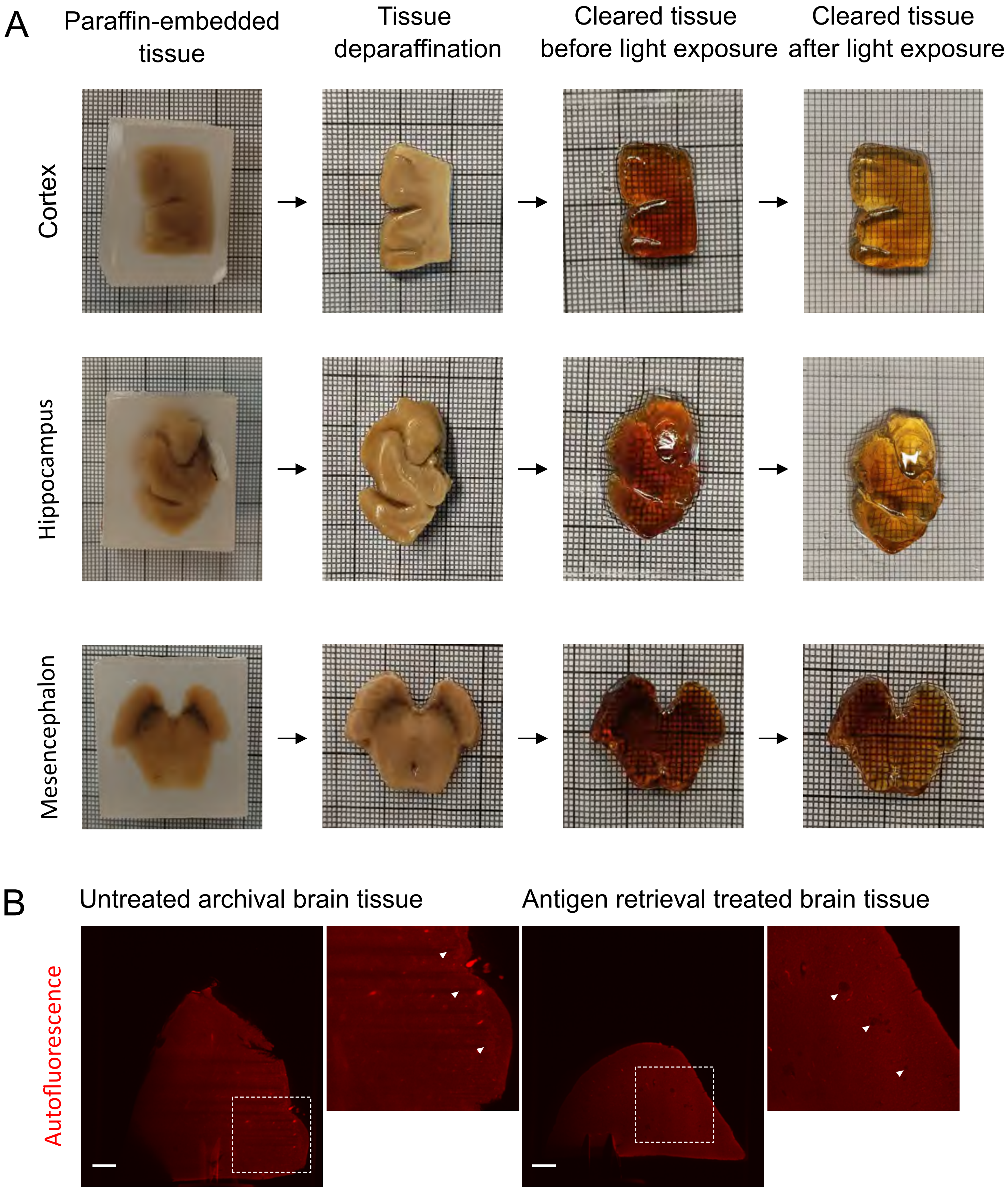


**Supplementary Figure 11 – Processing of archival brain tissue with CleLight. A)** Paraffin removal was achieved for different parts of the nervous system including cortex, hippocampus and mesencephalon (1mm per square). These samples could be then processed and cleared using CleLight. **B)** Comparison of untreated archival brain tissue containing PFA crystals (arrow heads) and its removal after a treatment with antigen retrieval. Scale bar= 1mm.

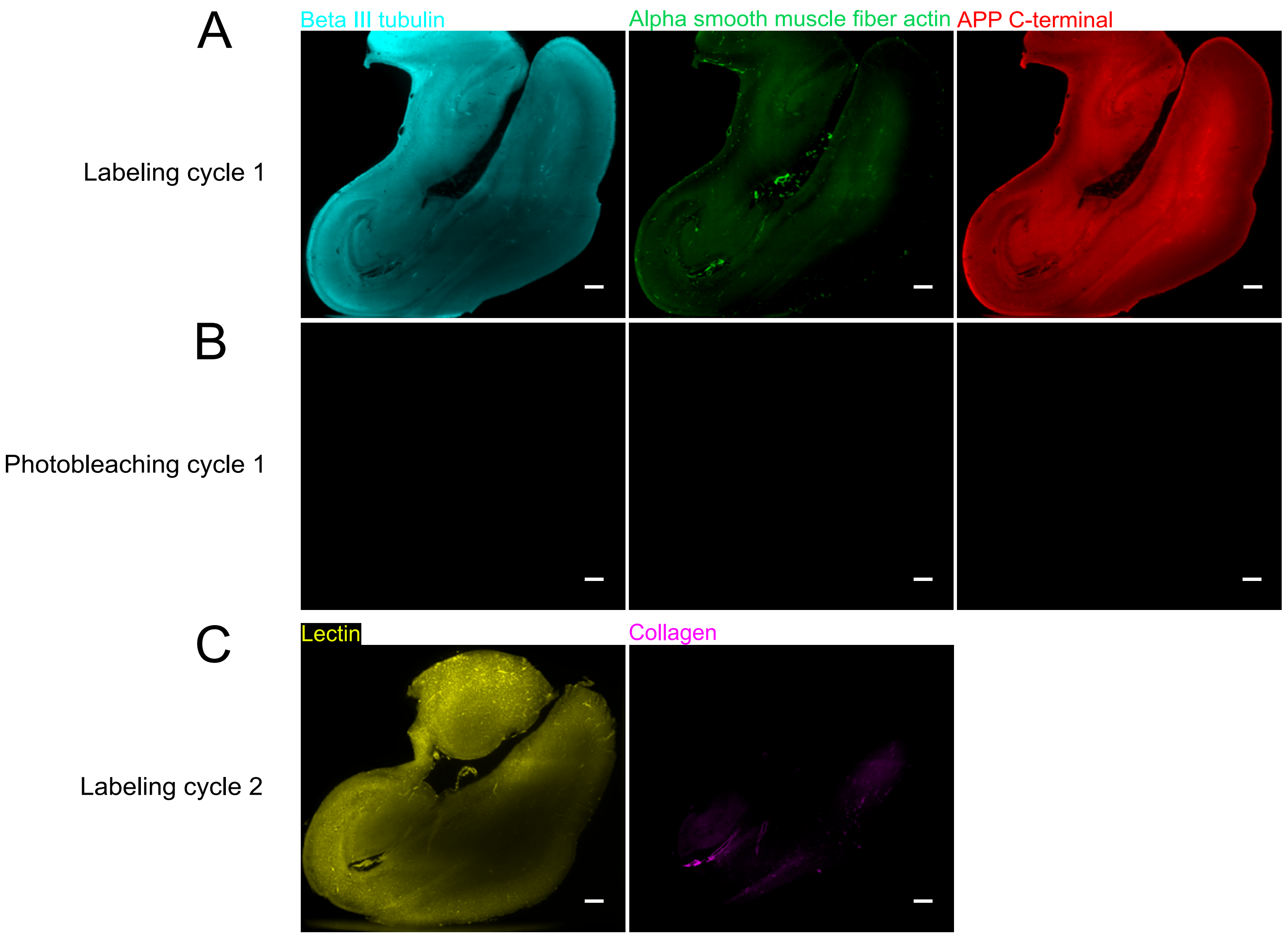


**Supplementary Figure 12 – Cyclic immunofluorescence with CleLight. Different rounds of labeling performed across the same hippocampus. A)** First round of labeling with alpha smooth muscle fiber actin (green), APP C-terminal (red) and Beta III tubulin (cyan). Scale bar= 1000µm. **B)** First round of photobleaching shows that fluorescence was completely quenched. Scale bar= 1000µm. **C)** Second round of labeling with lectin and collagen shows the preservation of the epitopes and of the tissue structure. Scale bar= 1000µm.

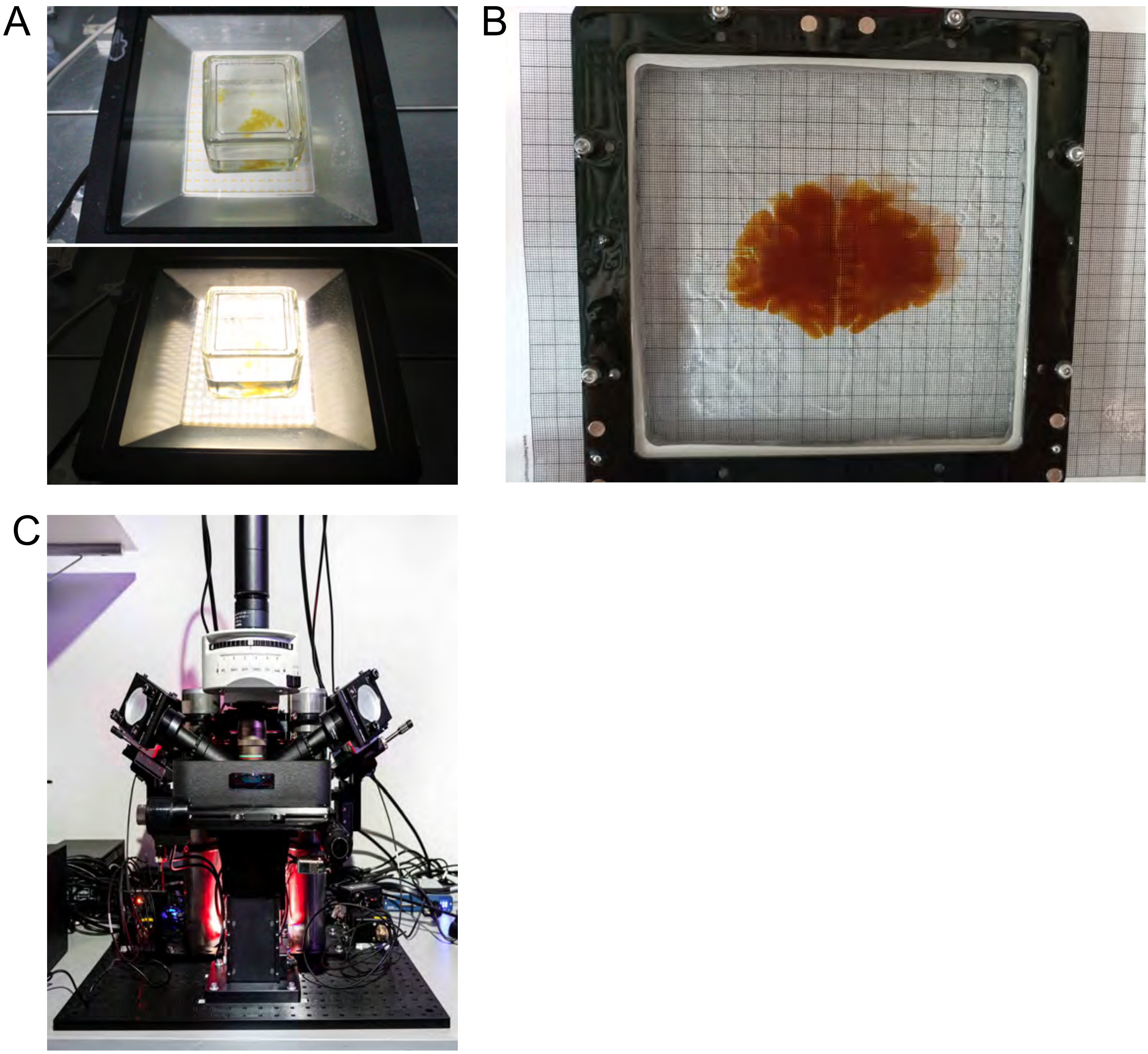


**Supplementary Figure 13 – Preparation of entire human brain slice for clearing and imaging. A)** Exposure of whole brain slice to white light. **B)** Sealed chamber containing DBE with an entire human brain slice before acquisition **C)** LSFM during acquisition of the slice.

**Supplementary tables**

**Supplementary Table 1** – Patient data for the samples used in this study.

| Case | Sex | Braak stage | Age of death | PMD (h) | Fixation time | Brain region used |
|---|---|---|---|---|---|---|
| Control 1 | M | - | 77 | 13 | 6 months | Frontal cortex |
| Control 2 | M | - | 70 | 15 | 2 years | Spinal cord<br>Cerebellum |
| Control 3 | M | - | 82 | 40 | 13 years | Substantia nigra |
| Control 4 | M | - | 77 | 38 | 13 years | Substantia nigra |
| Control 5 | M | - | 62 | 10 | 55 years | Frontal cortex |
| Alzheimer disease 1 | F | V-VI | 81 | 46 | 6 months | Frontal cortex |
| Alzheimer disease 2 | F | V-VI | 75 | 4 | 45 years | Frontal cortex |

**Supplementary Table 2** – Antibodies and chemical dyes used with CleLight.

| Antibody/Counterstain | Source and reference | Dilution | Secondary antibody |
|---|---|---|---|
| MAP2 | Synaptic Systems 188 004 | 1:250 | Alexa Fluor® 555 or 647 AffiniPure Fab Fragment Goat Anti-Guinea pig IgG (H+L). Jackson immunoresearch |
| Neurofilament axonal marker (SMI312) | BioLegend - 837904 | 1:250 | Alexa Fluor® 555 or 647 AffiniPure Fab Fragment Goat Anti-Mouse IgG (H+L). Jackson immunoresearch |
| Anti-beta III Tubulin | Abcam 18207 | 1:250 | Alexa Fluor® 555 or 647 AffiniPure Fab Fragment Goat Anti-Rabbit IgG (H+L). Jackson immunoresearch |
| Iba1 | FUJIFILM Wako Pure Chemical 019-19741 | 1:250 | Secondary antibodyCy™3 or 647 AffiniPure Donkey Anti-Rabbit IgG (H+L). Jackson immunoresearch |
| NeuN | Synaptic systems 266 004 | 1:250 | Alexa Fluor® 555 or 647 AffiniPure Fab Fragment Goat Anti-Guinea pig IgG (H+L). Jackson immunoresearch |
| Alexa Fluor® 647 Anti-NeuN antibody | Abcam 190565 | 1:250 | |
| Tyrosin hydroxylase | Merck millipore ab152 | 1:500 | Alexa Fluor® 555 or 647 AffiniPure Fab Fragment Goat Anti-Rabbit IgG (H+L). Jackson immunoresearch |
| VAMP2 | Synaptic Systems 104 211 | 1:250 | Alexa Fluor® 555 or 647 AffiniPure Fab Fragment Goat Anti-Mouse IgG (H+L). Jackson immunoresearch |
| Abeta 1-16 | BioLegend SIG-39320 | 1:500 | Secondary antibodyCy™3 or 647AffiniPure Donkey Anti-Mouse IgG (H+L). Jackson immunoresearch |
| Amyloid precursor protein (APP C-ter) | BioLegend SIG-39150 | 1:250 | Alexa Fluor® 555 or 647 AffiniPure Fab Fragment Goat Anti-Mouse IgG (H+L). Jackson immunoresearch |
| P-Tau AT8 | Thermo Fisher Scientific MN1020 | 1:250 | Alexa Fluor® 555 AffiniPure Fab Fragment Goat Anti-Mouse IgG (H+L). Jackson immunoresearch |
| Alpha -smooth muscle actin | Abcam 21027 | 1:250 | Alexa Fluor® 555 AffiniPure Fab Fragment Donkey Anti-Goat IgG (H+L). Jackson immunoresearch |

| | | | |
|---|---|---|---|
| Lectin DyLight™ 488 | Vector laboratories | 1:100 | - |
| TO-PRO-3 Iodide | Invitrogen T3605 | 1:100 | - |
| BioTracker 488 | Merck millipore SCT120 | 1:100 | - |
| Congo Red | Merck millipore 101340 | 1% | - |
| Thioflavin S | Sigma-Aldrich T-1892 | 1% | - |
| Fast green FCF | Thermo Scientific CatA16520.22 | 1:5000 | - |
